# Accurate and Sensitive Quantitation of the Dynamic Heat Shock Proteome using Tandem Mass Tags

**DOI:** 10.1101/696641

**Authors:** Aaron J. Storey, Rebecca E. Hardman, Stephanie D. Byrum, Samuel G. Mackintosh, Rick D. Edmondson, Wayne P. Wahls, Alan J. Tackett, Jeffrey A. Lewis

## Abstract

Cells respond to environmental perturbations and insults through modulating protein abundance and function. However, the majority of studies have focused on changes in RNA abundance because quantitative transcriptomics has historically been more facile than quantitative proteomics. Modern Orbitrap mass spectrometers now provide sensitive and deep proteome coverage, allowing direct, global quantification of not only protein abundance, but also post-translational modifications (PTMs) that regulate protein activity. We implemented, and validated using the well-characterized heat shock response of budding yeast, a tandem mass tagging (TMT), triple-stage mass spectrometry (MS^3^) strategy to measure global changes in the proteome during the yeast heat shock response over nine timepoints. We report that basic pH, ultra-high performance liquid chromatography (UPLC) fractionation of tryptic peptides yields superfractions of minimal redundancy, a crucial requirement for deep coverage and quantification by subsequent LC-MS^3^. We quantified 2,275 proteins across 3 biological replicates, and found that differential expression peaked near 90 minutes following heat shock (with 868 differentially expressed proteins at 5% FDR). The sensitivity of the approach also allowed us to detect changes in the relative abundance of ubiquitination and phosphorylation PTMs over time. Remarkably, relative quantification of post-translationally modified peptides revealed striking evidence of regulation of the heat shock response by protein PTMs. These data demonstrate that the high precision of TMT-MS^3^ enables peptide-level quantification of samples, which can reveal important regulation of protein abundance and regulatory PTMs under various experimental conditions.

## Introduction

Cells employ a diverse array of regulatory strategies to maintain homeostasis in the face of environmental challenges. These regulatory strategies exist on a number of physiological timescales. For instance, rapid responses (i.e. seconds to minutes) necessitate changing the activity of proteins that are already present in the cell, and can include changing concentrations of regulatory metabolites (e.g. allosteric inducers or inhibitors) or covalent post-translational protein modifications (PTMs). Over longer time scales (i.e. minutes to hours), cells can respond to stressful environments by remodeling gene expression. For example, the model eukaryote *Saccharomyces cerevisiae* responds to stress by globally remodeling gene expression to shift translational capacity towards the production of stress defense proteins that increase cellular fitness in the face of stress.^1-3^

While changes in protein abundance and activites are the ultimate mediators of biological responses, the majority of studies have historically focused on transcriptional changes. This was largely due to technical challenges related to accurately quantifying protein abundance.^4^ Independent of sequence, nucleic acids are chemically homogeneous enough to allow for the common hybridization chemistry used for microarrays,^5^ and subsequently for high-throughput sequencing.^6^ In contrast, proteins are much more heterogeneous and diverse in their chemistries, limiting our ability to design “one-size fits all” approaches to proteomics. Furthermore, it was generally assumed that while imperfect, mRNA levels were an adequate proxy for protein level estimates. However, early studies comparing the transcriptome and proteome in multiple organisms showed extremely poor correlations between mRNA and protein levels,^7-10^ creating a real doubt about whether mRNA levels were really a good proxy for protein levels. Some of this discrepancy may be due to experimental noise,^11^ highlighting both the challenge and importance of being able to accurately quantitate peptide abundance. However, while the correlation between mRNA and protein levels is not as poor as initially thought,^11, 12^ substantial regulation of protein levels still occurs post-transcriptionally. For example, detailed comparisons between mRNA and protein levels during the yeast response to hyperosmotic shock revealed that ∼80% of the variation in induced proteins can be explained by changes in mRNA abundance, with the remaining variation possibly explained by translational or post-translational regulation.^3^ Similarly, ribosomal profiling experiments have identified widespread changes in mRNA translational efficiency under a number of conditions ranging from meiosis to stress ^13-15^. Finally, it is well established that protein stability and degradation plays an important role in regulated protein turnover during environmental shifts.^16, 17^

In addition to protein abundance changes, PTMs are well known to regulate protein activity and/or stability during environmental shifts. For example, stress-activated protein kinases coordinate phospho-signal transduction cascades that are largely conserved from yeast through humans.^18-21^ Acetylation and methylation of histones and transcription factors also facilitate transcriptional reprogramming during stress.^22-26^ During stress, damaged proteins are frequently targeted for proteasomal degradation via ubiquitination.^16, 27^ Additionally, global changes in SUMOylation play an important role in heat stress adaptation,^28, 29^ providing further support for the notion that global PTM remodeling is a broad regulatory strategy during stress adaptation. Thus, an integrated view of stress physiology requires the ability to sensitively and accurately measure relative protein abundance and PTMs.

Here, we describe a workflow for using a tandem mass tagging (TMT) strategy to measure global proteomic changes during environmental shifts. The key advantage is that this approach supports simultaneous analyses of multiple samples in the same MS run. As proof-of-concept, we measured protein abundance changes in yeast before and after heat shock. The response to elevated temperature is arguably the best-studied environmental response across diverse organisms, with many important evolutionary features conserved from bacteria to humans.^30, 31^ Heat stress affects a number of cellular targets including increasing membrane fluidity^32^ (which leads to disruption of nutrient uptake,^33^ pH balance,^34, 35^ and ROS production due to “leaking” of electrons from the mitochondrial electron transport chain^36^), and protein unfolding (leading to induction of heat shock protein (HSP) chaperones and other protectants such as trehalose^37-39^).

While many studies have characterized the global transcriptional response to heat shock in yeast,^1, 2, 40-43^ relatively few studies have examined proteomic changes.^44, 45^ These previous studies have used either stable isotope labeling of amino acids in cell culture (SILAC) or label-free approaches, but to date no study has compared these methods to TMT, which has a potential advantage for multiplexing. We used a MultiNotch triple-stage mass spectrometry (MS^3^) workflow to quantify peptides, which mitigates interference of nearly isobaric contaminant ions that cause an underestimate of differential expression.^46, 47^ We tested our worklow using a 10-plex design comparing the proteome of unstressed cells to heat shocked cells over 9 timepoints, with three biological replicates.

Our TMT-MS^3^ workflow, which included pre-fractionation of peptide mixtures to reduce sample complexity and increase coverage of identifications, quantified over 2000 proteins between a heat shock and unstressed control sample. In addition to providing insight into the dynamics of protein abundance changes during heat shock, we also identified post-translationally modified peptides whose relative abundance also changed dynamically. These data demonstrate that the high precision of TMT-MS^3^ enables peptide-level quantification of samples, which can reveal important regulation of protein post-translational modifications under various experimental conditions.

## Experimental Procedures

### Sample Preparation for TMT-Labeling and LC-MS/MS

#### Yeast Growth Conditions

We first performed a pilot TMT 2-plex experiment comparing unstressed cells to heat stressed cells at a single timepoint, followed by a TMT-10 heat shock timecourse that was performed with three independent biological replicates. For both experiments, yeast strain BY4741 (S288c background; MATa *his3*Δ*1 leu2*Δ*0 met15*Δ*0 ura3*Δ*0*) was grown >7 generations to mid-exponential phase (OD_600_ of 0.3 to 0.6) at 25°C and 270 rpm in 500 ml of YPD medium (1% yeast extract, 2% peptone, 2% dextrose). For the TMT-2 experiment, cells were collected by centrifugation at 1,500 × *g* for 3 minutes, resuspended in either pre-warmed 25°C medium (unstressed sample) or 37°C medium (heat shocked sample), and then incubated for 1 hour at 25°C or 37°C, respectively. Following incubation, samples were collected by centrifugation at 1500 × *g* for 3 minutes, the media was decanted, and the pellet was flash frozen in liquid nitrogen and stored at −80°C until processing. For the TMT-10 experiment, heat shock was performed by adding an equal amount of 55°C preheated media, and cells were collected on cellulose nirate filters by vacuum filtration at 5, 10, 15, 30, 45, 60, 90, 120, and 240 minutes post-heat shock. Cells were immediately scraped from the filters into liquid nitrogen and stored at −80°C until processing.

#### TMT 2-Plex Sample Preparation

For the TMT-2 experiment, cell pellets were resuspended in 3 ml of lysis buffer (20 mM HEPES, 150 mM potassium acetate, 2 mM magnesium acetate, pH 7.4) plus EDTA-free Protease Inhibitor Tablets (Pierce catalog number 88266), and frozen dropwise into liquid nitrogen. Samples were lysed by cryogrinding using a Retsch MM 400 Mixer Mill (5 cycles of 30 Hz for 3 minutes), returning the chamber to liquid nitrogen between rounds. Proteins were thawed in cold water and precipitated with 4 volumes of ice cold acetone overnight, and resuspended in 5 ml of buffer containing 8 M urea, 5 mM dithiothreitol, and 1 M ammonium bicarbonate, pH 8.0. Ammonium bicarbonate was included to reduce protein carbamylation that occurs in urea-containing buffers.^48^ Samples were divided into 1 ml aliquots, flash frozen in liquid nitrogen, and stored at −80°C until further processing.

Protein samples were reduced by incubating with 5 mM tris(2-carboxyethyl)phosphine (TCEP) at 37°C for 1 hour, and alkylated with 15 mM iodoacetamide at room temperature for 30 minutes. Protein samples were extracted with chloroform-methanol, ^49^ resuspended in 100 mM triethylammonium bicarbonate (TEAB) pH 8.0 with 1 µg Trypsin per 50 µg protein, and incubated at 37°C for 16 hours. Tryptic peptides were desalted with Sep Pak C18 columns (Waters) according to the manufacturer’s instructions and lyophilized. Peptides were resuspended in 100 mM TEAB pH 8.0, quantified using a Pierce Quantitative Colorimetric Peptide Assay Kit (ThermoFisher Scientific), and equal amounds of peptide samples were labeled with Tandem Mass Tag (TMT) 2-plex reagents (ThermoFisher Scientific) according to the manufacturer’s instructions.

#### TMT 10-Plex Sample Preparation

For the TMT-10 experiment, cell pellets were resuspended in fresh 6 M guanidine HCl, 100 mM Tris-HCl, pH 8.0, and cells were lysed by incubation at 100°C for 5 minutes, 25°C for 5 minutes, and 100°C for 5 minutes. Proteins were precipitated by adding 9 volumes of 100% methanol, vortexing, and centrifuging at 9,000 x *g* for 5 minutes. The supernatant was carefully decanted, the protein pellets were air dried for 5 minutes, and then resuspended in 8 M urea.

Protein samples were diluted to 2M urea with 100mM Tris pH 8.0 and digested with a 1:50 ratio of Trypsin overnight at 25°C with gently mixing in the presence of 2.5mM TCEP and 10mM chloroacetamide. Digestion was quenched with 0.6% TFA to a pH less than 2 and peptides were desalted and lyophilized. Peptides were resuspended in 200mM TEAB to a final concentration of ∼8µg, quantitated with Pierce Colorimetric Peptide Assay, and diluted to 5µg/µl in TEAB. 100µl of each sample was mixed with a TMT label reconstituted in 50µl acetonitrile. Samples were incubated at RT for 1 hour. Labelling was quenched with 8µl 5% hydroxylamine for 15 minutes and samples were combined, desalted, and lyophilized.

### LC-MS/MS Data Analysis

For both TMT-2 and TMT-10, peptides were fractionated on a 100 mm × 1.0 mm Acquity BEH C18 column (Waters) using an UltiMate 3000 UHPLC system (ThermoFisher Scientific) with a 40-min gradient from 99:1 to 60:40 buffer A:B ratio under basic pH conditions (buffer A = 0.05% acetonitrile, 10 mM NH_4_OH; buffer B = 100% acetonitrile, 10 mM NH_4_OH). The individual fractions were then consolidated into 24 super-fractions.

Super-fractions from the TMT 2-plex experiment were loaded on a Jupiter Proteo resin (Phenomenex) on an in-line 150 mm × 0.075 mm column using a nanoAcquity UPLC system (Waters). Peptides were eluted using a 45-min gradient from 97:3 to 65:35 buffer A:B ratio (buffer A = 0.1% formic acid; buffer B = acetonitrile, 0.1% formic acid) into an Orbitrap Fusion Tribrid mass spectrometer (ThermoFisher Scientific). MS acquisition consisted of a full MS scan at 240,000 resolution in profile mode of scan range 375-1500, maximum injection time of 400 ms, and AGC target of 5 × 10^5^, followed by CID MS/MS scans of the N most abundant ions of charge state +2-7 within a 3 second duty cycle. Precursor ions were isolated with a 2 Th isolation window in the quadrupole, fragmented with CID at 35%, and analyzed in the ion trap with a maximum injection time of 35 ms and a scan setting of Rapid. Dynamic exclusion was set to 20 seconds with a 10 ppm tolerance. MS^2^ scans were followed by synchronous precursor selection and HCD (65%) fragmentation of the 10 most abundant fragment ions. MS^3^ scans were performed at 30,000 resolution with a maximum injection time of 200 ms and AGC target of 100,000.

For the TMT 10-plex experiment, super fractions were loaded on a 150 mm × 0.075 mm column packed with Waters C18 CSH resin. Peptides were eluted using a 45-min gradient from 96:4 to 75:25 buffer A:B ratio into an Orbitrap Fusion Lumos mass spectrometer (ThermoFischer Scientific). MS acquisition consisted of a full scan at 120,000 resolution, maximum injection time of 50 ms, and AGC target of 7.5 × 10^5^. Selection filters consisted of monoisotopic peak determination, charge state 2-7, intensity threshold of 2.0 × 10^4^, and mass range of 400-1200 m/z. Dynamic exclusion length was set to 15 seconds. Data-dependent cycle time was set for 2.5 seconds. Selected precursors were fragmented using CID 35% with an AGC target of 5.0 × 10^3^ and a maximum injection time of 50 ms. MS^2^ scans were followed by synchronous precursor selection of the 10 most abundant fragment ions, which were fragmented with HCD 65% and scanned in the Orbitrap at 50,000 resolution, AGC target of 5.0 × 10^4^ and maximum injection time of 86 ms.

Proteins were identified by database search using MaxQuant^50^ (Max Planck Institute) using the Uniprot *S. cerevisiae* database from October 2014,^51^ with a parent ion tolerance of 3 ppm and a fragment ion tolerance of 0.5 Da. Carbamidomethylation of cysteine residues was used as a fixed modification. Acetylation of protein N-termini and oxidation of methionine were selected as variable modifications. Mascot searches were performed using the same parameters as above, but with peptide N-terminal fixed modification of TMT 2-plex (+225.16) or TMT 10-plex (+229.16), and variable modifications of diglycine (+114.04) and TMT 2- or 10-plex on lysine residues, and phosphorylation (+79.97) of serine and threonine. Mascot search results were imported into Scaffold software (v4)^52^ and filtered for protein and peptide false discovery rates of 1%. Data normalization and analyses were performed using R, and all R scripts for analysis are provided in File S1. Reporter ion intensities were log_2_ transformed, mean-centered for each spectrum, then median-centered for each channel. Peptide and protein quantitative values were obtained by taking the average of the quantitative values for all spectra mapping to the peptide or protein. All mass spectromentry data and MaxQuant search results have been deposited to the ProteomeXchange Consortium via the PRIDE^53^ partner repository with the dataset identifier PXD014552 and 10.6019/PXD014552.

Proteins with significant abundance differences in response to heat at each time point relative to the unstressed control were identified by performing an empirical Bayes moderated *t-* test using the BioConductor package Limma v 3.36.2 and Benjamini-Hochberg FDR correction.^54^ Unless otherwise stated, we applied an FDR cutoff of 0.05 (see File S2 for the Limma output). Protein or peptide clustering was performed with Cluster 3.0 (http://bonsai.hgc.jp/~mdehoon/software/cluster/software.htm) using hierarchical clustering and Euclidian distance as the metric.^55^ Timepoints were weighted using a cutoff value of 0.4 and an exponent value of 1. Functional enrichments of gene ontology (GO) categories were performed using GO-TermFinder (https://go.princeton.edu/cgi-bin/GOTermFinder),^56^ with Bonferroni-corrected *P*-values < 0.01 taken as significant. Complete lists of enriched categories can be found in File S3. Significantly enriched regulatory associations were identified using the YEAst Search for Transcriptional Regulators And Consensus Tracking (YEASTRACT) database,^57^ using documented DNA binding plus expression evidence. Significant associations can be found in File S4.

### Quantitative Western Blotting

Validation of LC-MS/MS was performed using the yeast TAP Tagged ORF collection (GE Dharmacon) in biological triplicate. Cells were collected and heat shocked exactly as described for the TMT-2 sample preparation, with the duration of the 37°C heat shock being 1 hour. The OD_600_ for the heat-shocked and unstressed control samples were recorded for subsequent normalization. Fifteen ml of each sample was collected by centrifugation at 1500 x *g* for 3 minutes, the media was decanted, and the pellet was flash frozen in liquid nitrogen and stored at −80°C until processing. Sample processing for western blotting was performed as described^58^ with the following modifications. Samples were thawed and re-suspended in 1 ml of sterile water, and then an equal number of cells (∼ 1 × 10^7^) was removed and collected by centrifugation at 10,000 x *g* for 1 minute. Cells were resuspended in 200 µl lysis buffer (0.1 M NaOH, 50 mM EDTA, 2% SDS, 2% β-mercaptoethanol plus EDTA-free Protease Inhibitor Tablets (Pierce catalog number 88265)). Samples were then incubated at 90°C for 10 minutes, 5 µl of 4 M acetic acid was added, and the sample was vortexed at maximum speed for 30 seconds. Samples were then incubated at 90°C for an additional 10 minutes to complete lysis. Fifty µl of loading buffer (0.25 M Tris-HCl pH 6.8, 50% glycerol, 0.05% bromophenol blue) was added to each sample, and samples were centrifuged at 21,130 × *g* for 5 minutes to pellet cellular debris. Twenty µl of each sample was loaded onto a 4 - 20% gradient acrylamide gel (Bio-Rad), separated by SDS-PAGE, and transferred for 1 hour onto an Amersham Protran Premium 0.45 nitrocellulose membrane (GE Healthcare). Western blots were performed using a mixture of mouse anti-actin antibodies (VWR catalog number 89500-294) and rabbit anti-TAP antibodies (ThermoFisher catalog number CAB1001) to simultaneously detect Act1p and the Tap-tagged protein of interest. Anti-TAP and anti-actin antibodies were used at a dilution of 1:1000 and 1:2500, respectively. IRDye^®^ 680RD-conjugated anti-rabbit IgG (LI-COR Biosciences catalog number 926-68073) and IRDye^®^ 800CW-conjugated anti-mouse IgG (LI-COR Biosciences catalog number 926-32212) were used as secondary antibodies at a dilution of 1:10,000. Detection was performed with a LI-COR Odyssey Imaging System using Image Studio v2.0. Densitometry was performed using ImageJ,^59^ and log_2_ fold changes upon heat shock were calculated for each TAP-tagged protein following normalization to actin. Raw images and pixel densities can be found in Files S5 and S6.

## Results and Discussion

### Precision of MultiNotch MS^3^

We first sought to measure the precision of our TMT-MS^3^ workflow by characterizing the yeast response to elevated temperatures. One of the great advantages of TMT is the ability to multiplex, though there are mixed reports concerning whether increased multiplexing comes at the expense of protein identification and/or accuracy.^60, 61^ Thus, we first compared the accuracy and total protein identification of TMT-2 plex vs. TMT-10 plex. For our pilot TMT-2 experiment, we measured changes in protein abundance before and after 60 minutes of a 25°C to 37°C heat shock. We then performed a TMT-10 plex experiment designed to capture proteome dynamics of cells responding to heat shock over 9 timepoints in biological triplicate (Figure 1).

**Figure 1.**
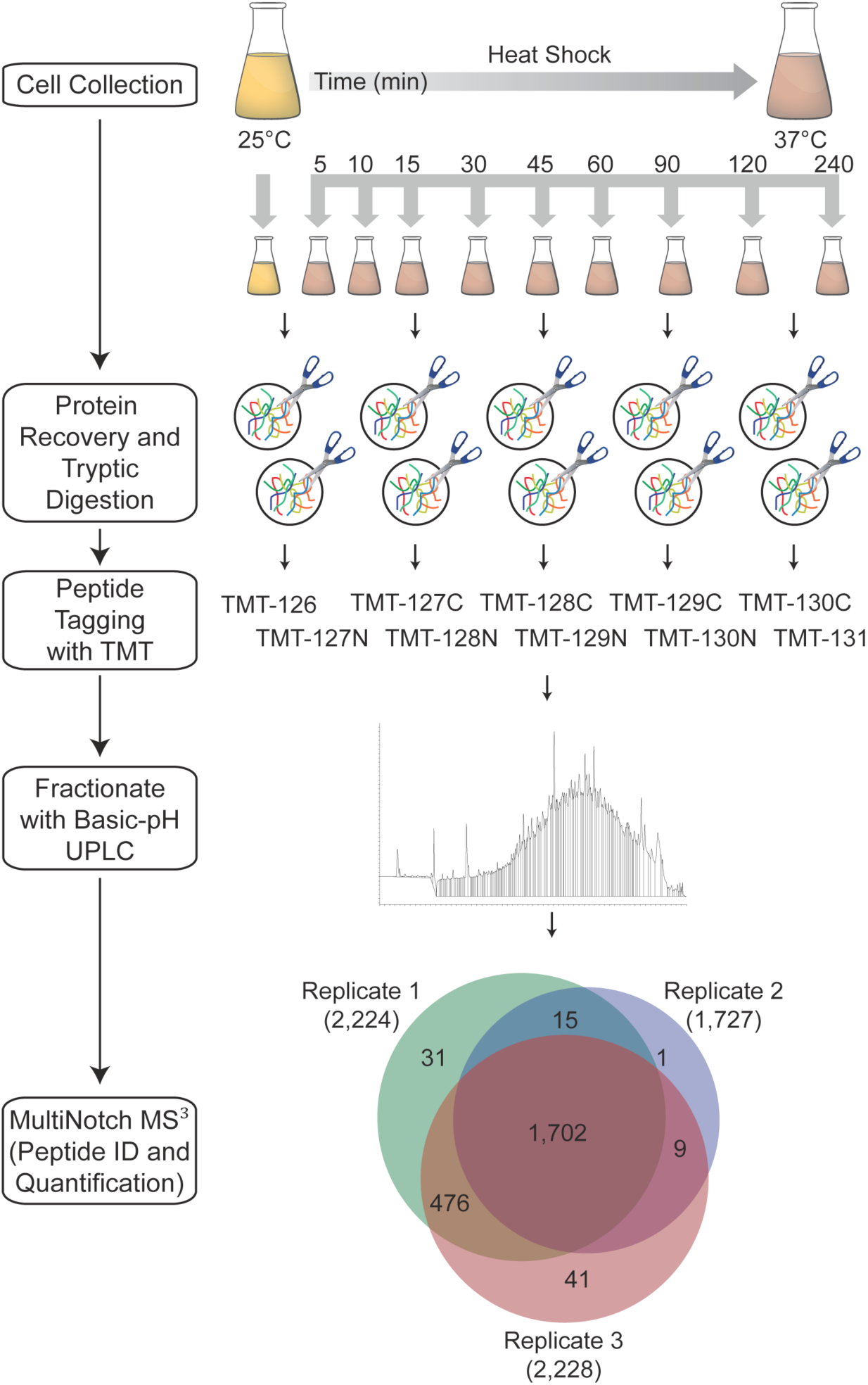
Schematic of Proteomic Workflow. Yeast cells were grown to mid-exponential phase at 25°C, an unstressed control sample was collected, and then cells were subjected to a 37°C heat shock, with samples collected at the indicated time points. Protein samples were digested with trypsin, labeled with one of the 10-plex tandem mass tags (TMT-10), pooled and fractionated by high-pH UPLC, and analyzed by MultiNotch LC-MS^3^. The Venn diagram depicts the overlap for proteins identified in each biological replicate.

For both sets of experiments, cells were harvested and lysed, and then protein samples were prepared using a standard bottom-up proteomics workflow with in-solution trypsin digestion, TMT 2-plex or 10-plex labeling, peptide fractionation, and analysis via MultiNotch MS^3^ on an Orbitrap Fusion (TMT-2) or Orbitrap Fusion Lumos (TMT-10).

One of the major challenges in quantitative proteomics is that peptides exist across a broad dynamic range of abundances, with high-abundance peptides dominating the signal of complex samples.^62^ Reducing sample complexity through fractionation improves the ability to detect and quantify low abundance peptides.^63^ In an ideally-resolved sample, peptides are found within a single fraction. One widely used workflow for offline sample fractionation is the separation of proteins by SDS-PAGE prior to trypsin digestion. However, this workflow is incompatible with TMT labeling, as proteins must be digested and labeled prior to fractionation. An alternative fractionation procedure is to separate peptide species by high-performance liquid chromatography (HPLC), with fractionation via basic-pH reversed-phase HPLC showing the best peptide coverage for complex human proteomes.^64, 65^

Thus, we fractionated our samples using basic-pH ultra-high performance liquid chromatography (UPLC) into 96 fractions, which were pooled into 24 superfractions. To measure the resolving power of basic-pH UPLC, we used the TMT 2-plex experiment to analyze the number of superfraction(s) in which each unique peptide was found (Figure 2A). By this analysis, 84% of peptides were found within a single superfraction, and 97% of peptides were found in two or fewer fractions. Additionally, peptides were evenly distributed across superfractions, with each superfraction yielding approximately 900 unique peptides (Figure 2B). Thus, basic-pH UPLC fractionation suitably reduces the amount of redundant MS^2^ and MS^3^ scans and increases the depth of unique peptide and protein identifications, which is important because the MultiNotch MS^3^ method has a slightly slower duty cycle than MS^2^-based reporter ion quantitation.

**Figure 2.**
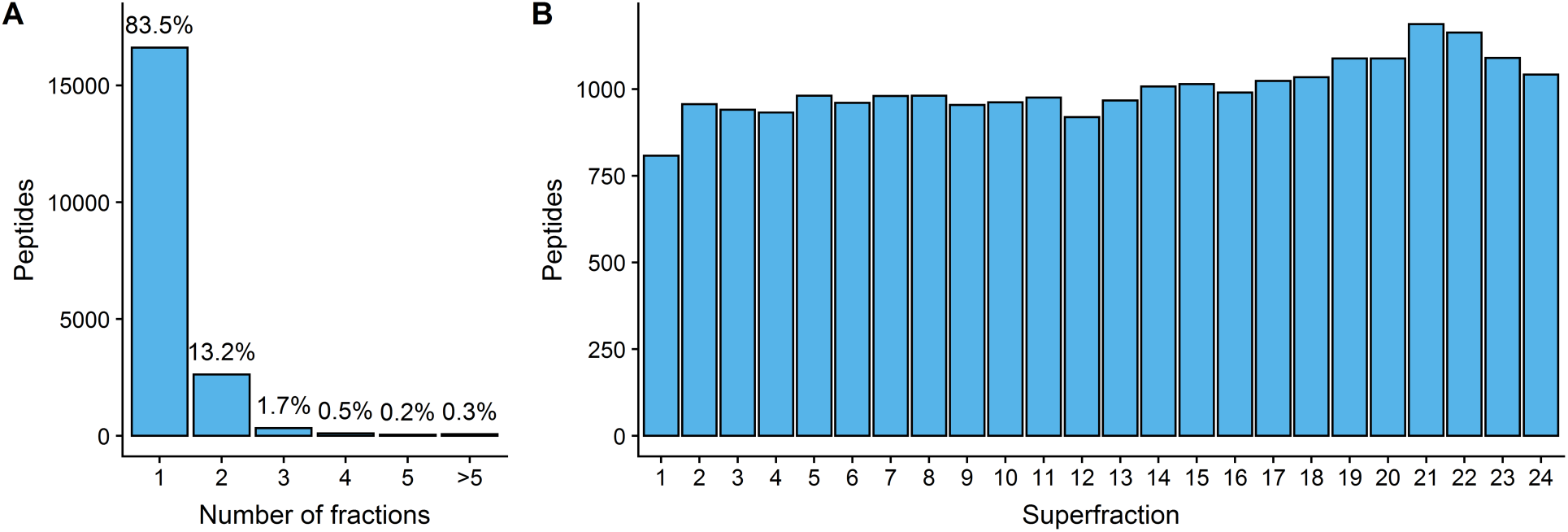
High pH UPLC fractionation efficiently separates TMT-labeled peptides. Protein samples were digested with trypsin, TMT 2-plex labeled, and then fractionated by UPLC under basic conditions, and the fractions were concatenated into 24 superfractions. Each superfraction was then analyzed by LC-MS^3^. **A**) Resolution of high pH fractionation. The X-axis shows the number of superfractions in which a given peptide was detected (bins); the histogram (Y-axis) shows the number and frequency of peptides in each bin. Note that the vast majority of peptides partition within only one or two of the 24 superfractions, greatly reducing sample complexity and increasing depth of coverage by LC-MS^3^. **B**) Sampling depth for each fractionation. The bar graph shows the number of peptides identified (Y-axis) in each superfraction (X-axis).

We next tested the precision of the MultiNotch MS^3^ method. As a first measure of the precision of TMT proteomics, we used the TMT 2-plex experiment to analyze the coefficient of variation (CV) of log_2_ fold changes in response to heat stress for each unique peptide (7,129 total) identified by MaxQuant (Figure 3A). The CV values for 85% of quantified peptides were below 30% (6070/7129). The distribution of both standard deviations and CV% were lowest for peptides with two spectral counts, which represented the largest class of peptides (4759/7129). This was somewhat unexpected, as standard deviation is generally inversely proportional to sample size (or peptide abundance in our case). Importantly, this trend was also observed across three independent TMT 10-plex experiments (Figure S1). Additionally, we identified a larger number of unique peptides (22696, 12025, 36080 for Replicates 1-3) and similar number of total proteins (2229, 1731, 2233 for Replicates 1-3) in the TMT-10 experiment, suggesting that both high accuracy and multiplexing are achievable without a large tradeoff in peptide identification. Overall, the data suggests that our TMT proteomic workflow yields reliable measurements for low abundance proteins, which we sought to examine in more detail.

**Figure 3.**
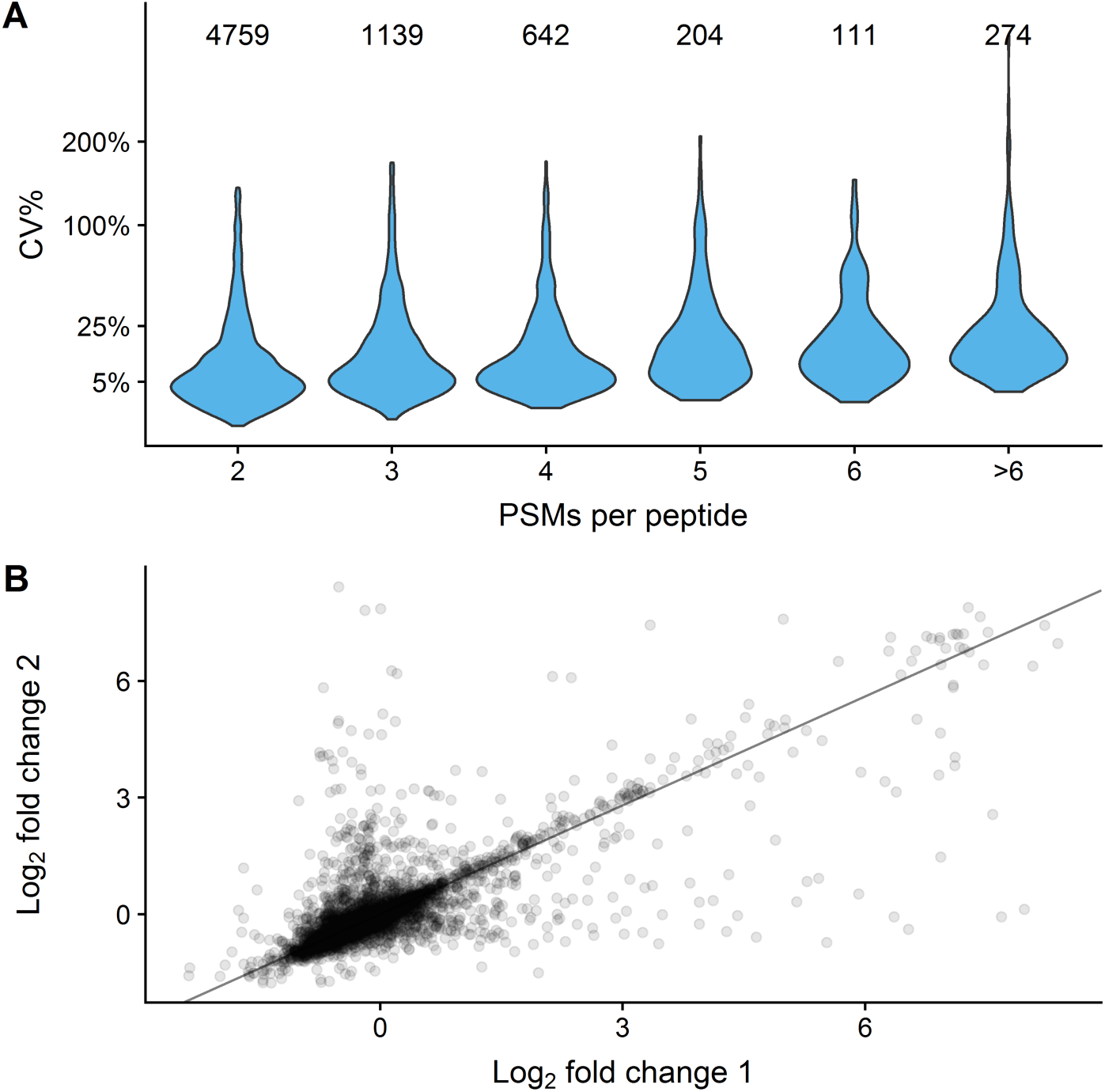
TMT-MS^3^ measurements of peptides are highly precise, even for low-abundance species. An underlying principle of quantitative proteomics is that high-abundance proteins yield more peptide spectral counts or measurements (PSMs) than low-abundance proteins. The differential abundance of peptides (fold change) following 60 minutes heat shock was used to determine the precision of PSM measurements for TMT-2. **A**) Precision of measurements as a function of PSM number. Violin plots showing coefficient of variation (CV) of fold change measurements for peptides with different numbers of PSMs. The number above each plot shows the number of data points in each plot. **B)** Precision of measurements for low-abundance peptides. Scatter plot shows log_2_ fold changes for 4,759 peptides with two spectral counts.

To further examine the precision of low-abundance peptides in the TMT 2-plex dataset, we filtered the data for peptides with 2 spectral counts, and analyzed the precision of the two peptide measurements. The data followed a right-skewed distribution, with the majority (95%) of peptides showing good agreement between measurements (*r*^*2*^ = 0.87, Figure 3B). Including the values from the remaining 5% of peptides markedly decreases the goodness-of-fit for all data points (*r*^*2*^ = 0.54), indicating that a small frequency of outlier measurements pose a challenge in this workflow. However, we conclude that the MultiNotch method yields high-quality data for the vast majority (95%) of peptide-level measurements, and there remains a need to predict the quality of a measurement in the absence of technical replicates, possibly through measuring the proportion of SPS ion intensity which did not map to the matched peptide.

### Using Increased Multiplexing to Characterize the Dynamic Heat Shock Proteome

To determine whether our TMT workflow was able to recapitulate known biology while providing new insights, we examined the dynamic response to heat stress across 9 timepoints (from 5 – 240 minutes post heat shock) using TMT 10-plex reagents and three independent biological replicates. Out of 2,276 proteins with at least duplicate data, 1,148 proteins were differentially expressed (FDR < 0.05) in at least one time point. We used quantitative western blotting on 7 signficantly induced proteins at the 60-minute time point, and all 7 proteins independently validated the proteomic data (Figure S2). We also compared our data to two yeast heat shock proteomic studies. First, we compared our dataset to a SILAC study from Nagaraj et al. that looked at a 30 minute heat shock.^44^ Compared to the SILAC experiment, we identified fewer differentially expressed proteins at 30 minutes post heat shock (150 vs. 234, Figure S3). This is likely due to a larger number of proteins identified by Nararaj et al. (3,152 vs. 2,276) combined with more statistical power due to an additional biological replicate. Notably, the proportion of proteins identified as significantly differentially expressed was similar across both studies (7.4% in Nagaraj et al. vs. 6.6% in this study). We next compared our data to a recent label-free study from Jarnuczak et al. that measured the heat shock response over 5 time points,^45^ specifically their time point with the highest number of differentially expressed proteins (240 minutes). While their study had more statistical power due to an additional biological replicate, we still identified more proteins as differentially expressed (Figure S3), and at a higher proportion (20.1% in Jarnuczak et al. vs. 27.0% in this study). Some of these differences in the ability to identify differentially expressed proteins are likely due to differences in experimental design (e.g. choice of heat shock temperature). However, some of the differences are likely due to the increased precision and lower variance of TMT-MS^3^, as smaller fold changes were more likely to be called significant in our dataset.

Examining the the most up- and down-regulated processes (>1.5-fold) during heat shock revealed processes likely important for acclimation to elevated temperatures. Proteins with significantly higher expression (>1.5-fold) following heat shock were enriched for functions known to be important for tolerating elevated temperatures. These included processes related to protein refolding (*p* = 3 × 10^−14^), oxidative stress response (*p* = 3 × 10^−5^), and metabolism of the known stress protectant molecule trehalose (*p* = 3 × 10^−3^). Other metabolic processes were also induced, including those related to redox chemistry (*p* = 3 × 10^−21^), amino acid metabolism (*p* = 2 × 10^−6^), and nucleotide metabolism (*p* = 5 × 10^−6^). In contrast, proteins repressed during heat shock were enriched for functions related to ribosomal biogenesis (*p* = 1 × 10^−22^), RNA processing (*p* = 8 × 10^−14^), and gene expression (*p* = 2 × 10^−7^).

We next sought to take advantage of our time series data to analyze the dynamics of the heat shock response in more detail. The maximal response occurred between 60 and 90 minutes post heat shock based on both the number of differentially expressed genes and the magnitude of the changes (Figures 4 and 5C). We identified 10 proteins with significantly higher abundance within 10 minutes post heat shock that likely reflect increased stabilization of key proteins. Of these 10 proteins with extremely early “induction,” four are heat shock protein chaperones (Hsp26p, Hsp42p, Hsp78p, Hsp104p), two are ribosomal proteins (Rps29bp, Rpl35ap), two are metabolic enzymes (Ura1p, Gre3p), and two are involved in protein targeting (Btn2p, Vac8). Intriguingly, the protein sorting protein Btn2p—the only protein with significantly increased abundance at the 5 min time point—works with the chaperone Hsp42p to regulate compartmentalization of protein aggregates for later repair or removal.^66^ Btn2p is known to be rapidly degraded by the proteasome during unstressed conditions,^67^ suggesting that changes in protein stability may play an important role in the early stages of the heat shock response.

**Figure 4.**
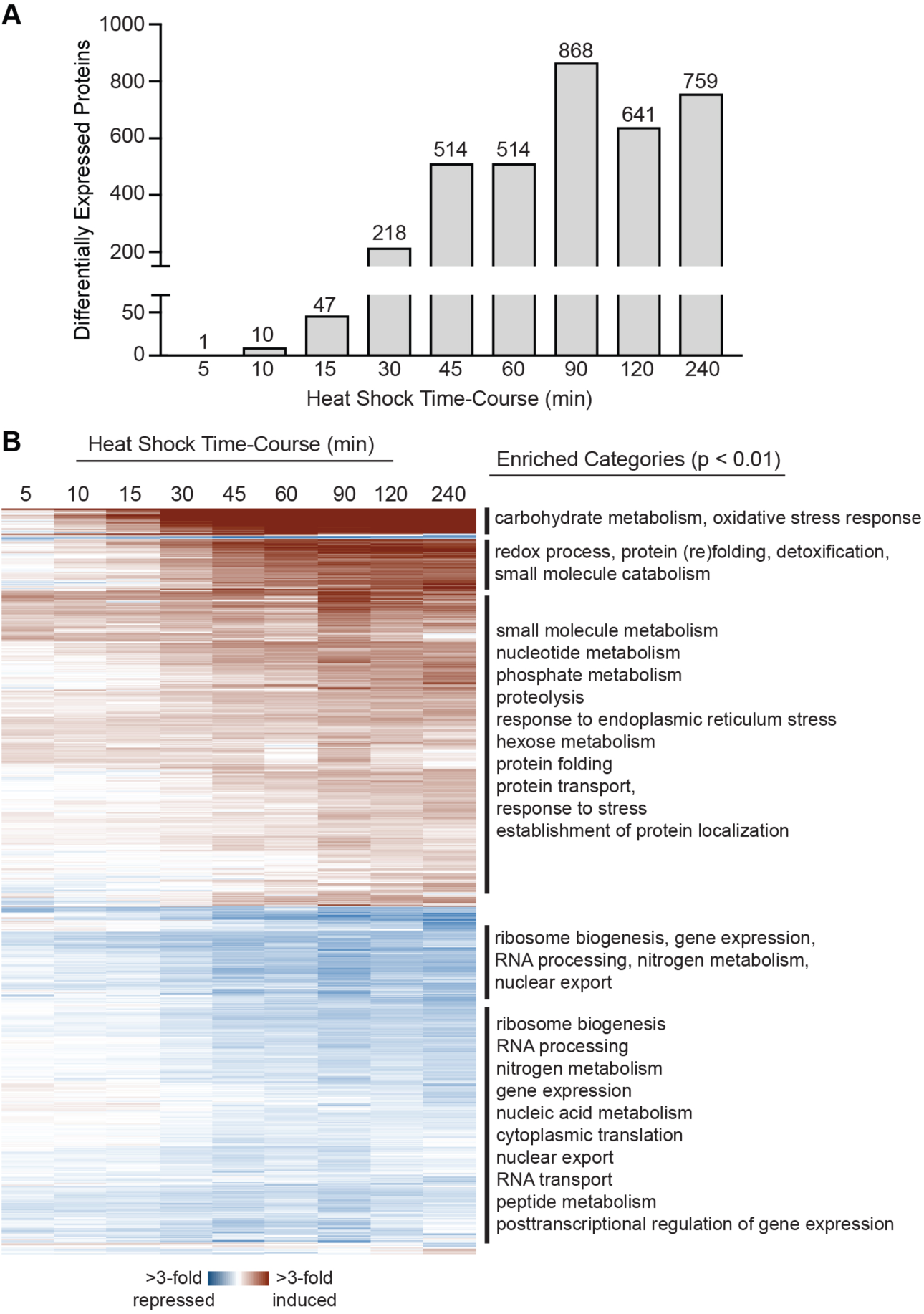
The temporal dynamics of the heat shock proteome. **A)** The number of differentially expressed proteins (FDR < 0.05) for each time point. **B)** The heat map depicts hierarchical clustering of 1,148 proteins whose change in abundance was statistically significant (FDR < 0.05) at any time point. Each row represents a unique protein, and each column represents the average expression change of heat stressed vs. unstressed cells at each time point. Red indicates induced and blue indicates repressed expression in response to heat. Enriched functional groups (Bonferroni-corrected *P* < 0.01, see Methods) are annotated to the right. Complete Gene Ontology (GO) enrichments for each cluster can be found in File S3.

**Figure 5.**
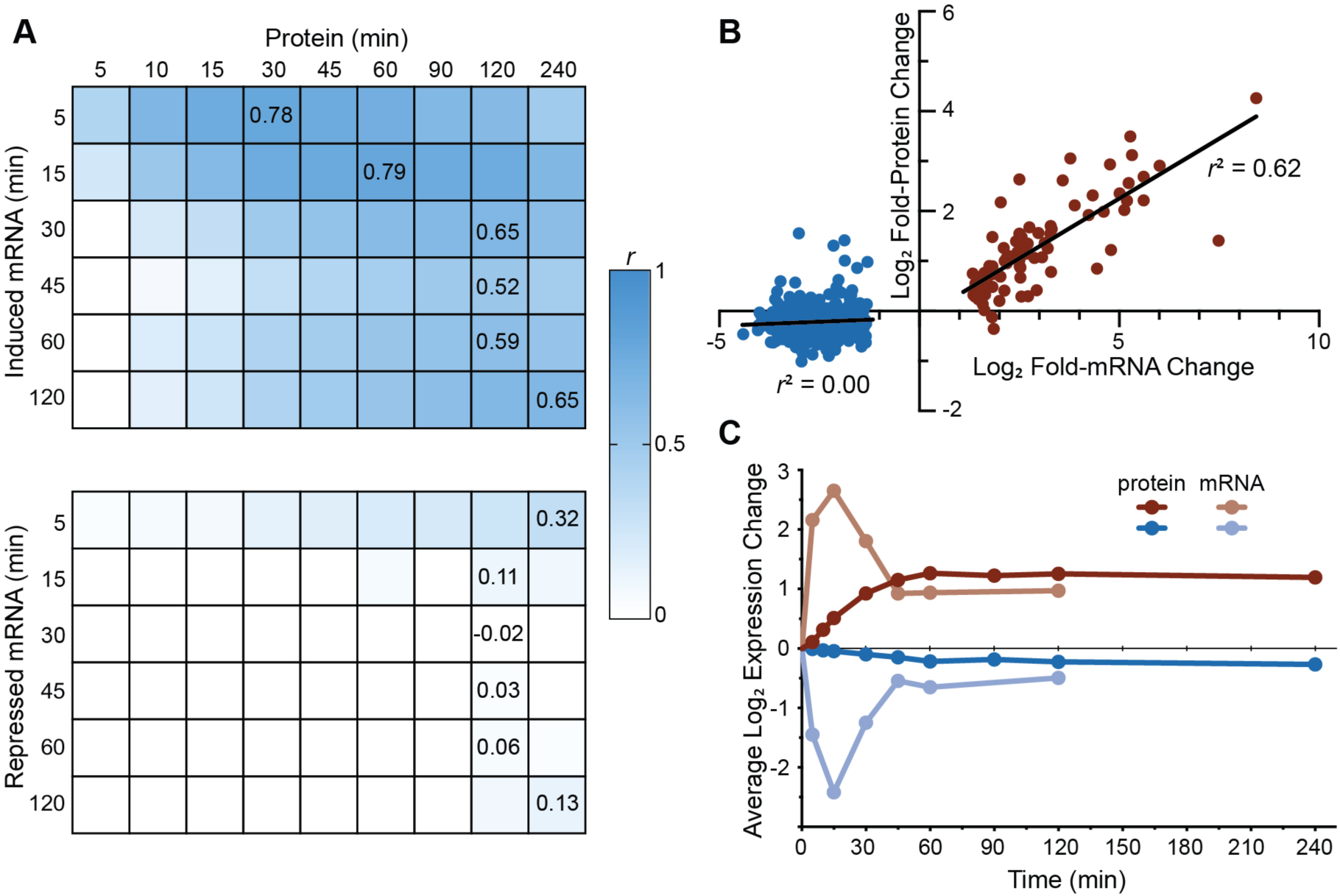
Heat shock responsive mRNAs correlate with protein induction but not repression. **A**) The tables compare changes in protein abundance at each time point following heat shock to mRNAs that significantly increased (top) or decreased (bottom) in expression reported by Eng et al.^40^ (at FDR < 0.05). Shading indicates Pearson correlation coefficients (r) between the the two datasets, with values of maximal concordance reported. **B)** Plot of the correlation between induced (red) and repressed (blue) mRNAs at 15 minutes vs. their corresponding proteins at 60 minutes. **C)** The temporal dynamics for all significantly induced or repressed mRNAs and their corresponding proteins.

To better identify temporal patterns in our data, we hierarchically clustered the 1,148 proteins with differential expression (Figure 4B). Induced proteins could be roughly categorized into three clusters: rapid and strong responders (41 proteins with a peak response of 60 minutes, average log_2_ fold change of 2.33), moderately induced responders (81 proteins with a peak response of 90 minutes, average log_2_ fold change of 1.18), and a broad cluster of lowly induced responders (469 proteins with a 90-min peak response, average log_2_ fold change of 0.50). The 41 rapid responders included several proteins known to be involved in the first line of heat stress defense including several key chaperones (Hsp26p, Hsp42p, Hsp78p, Hsp104p), glycogen and trehalose metabolic enzymes (Glc3p, Gsy2p,Tsl1p, Gph1p), and aromatic amino acid catabolic enyzmes (Aro9p, Aro10p). Additionally, there were several induced aldehyde dehydrogenases (Ald2p, Ald3p, Ald4p) and proteins involved in carbohydrate metabolism (Glk1p, Hxk2p, Gre3p, Sol3p, Yjr096wp, Pgm2p). The second wave of moderate responders also included additional chaperones or co-chaperones (Aha1p, Cpr1p, Cpr3p, Cpr6p, Hsp60p, Hsp82p, Sis1p, Ssa3p, Ssa4p, Sti1p) and trehalose metabolic enzymes (Nth1p and Tps2p). The broad lowly-induced responders included heat shock chaperones (Hsc82p, Ssa1p, Ecm10p, Ssc1p, Ssa2p, Kar2p, Sse2p, Mdj1p), but was also enriched for diverse metabolic functions including nitrogen metabolism (*p* = 3 × 10^−14^), nucleotide metabolism (*p* = 5× 10^−14^), phosphorus metabolism (*p* = 1 × 10^−9^), and glucose metabolism (*p* = 4 × 10^−5^).

We hypothesized that the rapid responders may largely represent proteins that directly respond to heat shock, while the lowly induced responders may reflect indirect responses. To test this, we used the YEASTRACT database to identify transcription factors that may be regulating gene expression for each cluster. The two major transcription factors that regulate the heat shock response in yeast include the heat shock transcription factor Hsf1p,^68^ and the paralogous general stress-responsive transcription factors Msn2p and Msn4p.^69^ Genes encoding rapid responders were more likely to regulated by Hsf1p (83%) than either the moderately (59%) or lowly (36%) genes (File S4). Similarly, the rapid response genes were also more likely to be regulated by Msn2p/4p (95%) than the moderately (84%) and lowly (60%) induced genes. Additionally, we identified several transcription factors that regulate the lowly-expressed genes that are themselves either Msnp2/4p targets (Rdr1p, Xbp1p, Oaf1p) or both Msn2/4 and Hsf1 targets (Rap1p, Tup1p, Pho4p, Ino2p, Tye7p, Reb1p, Mth1p, Prs1p). These transcription factors regulate diverse processes that are likely indirectly impacted by heat shock, including cell cycle progression and lipid, glucose, and phosphate metabolism.

Lastly, we examined the relationship between mRNA induction during heat shock and the proteomic response. Jarnuczak et al. also measured the correlation of mRNA and protein level changes, and found a modest correlation (*r* = 0.49 for the pairwise comparision with the highest correlation). We performed a similar analysis using a heat shock microarray time course (5, 15, 30, 45, 60, and 120 minutes) from Eng et al.,^40^ and we found a stronger correlation between changes in the heat shock transcriptome and proteome (*r* = 0.71). We explored the relationship between mRNA and protein further, which revealed some fundmental differences between how mRNAs and proteins are regulated during heat shock. First, in contrast with mRNA expression—where more mRNAs are repressed than induced during heat shock—we found that more proteins had increasing rather than decreasing abundance changes. This likely reflects the fact that proteins are more stable than mRNAs and that targeted protein degradation may be necessary to rapidly decrease protein levels.^70, 71^ Similar to Lee and colleagues^3^, we found that changes in mRNA abundance for induced transcripts correlated rather well with protein induction, while protein abundances changes were showed poor correlation with repressed mRNAs. This is consistent with proposed models suggesting the function of transcript repression is not to reduce protein abundance for those transcripts, but instead frees ribosomes and increases translation of the induced transcripts.^3, 72^ Intriguingly, despite the apparent lack of correlation for repressed mRNAs and their corresponding proteins, we did find that the functional enrichments for repressed mRNAs and repressed proteins were similar (i.e. ribosome biogenesis and translation). The poor correlation of repressed mRNAs and proteins occurs largely because repressed mRNAs show a wide range of repression values, while repressed proteins largely cluster around 1.5-fold repression (Figure S4). This “buffering” of repressed protein-level changes could be due to the increased stability of proteins vs. mRNAs.^70^ The repressed proteins are strongly enriched for the pre-ribosome complex (*p* = 3 × 10^−37^), thus the buffering of repressed proteins towards similar relative levels may help maintain proper subunit stoichiometry during stress.

### Analysis of protein post-translational modifications

The proteomics community uses several different peptide search engines for peptide spectrum matching, with each search engine having various strengths and weaknesses. Software with high-performance PTM identification may not be compatible with TMT-MS^3^ quantitation, and software with TMT-MS^3^ quantitation may fail to identify modified peptides in a sample. For example, we have previously used Mascot Distiller to search for post-translational modifications,^73^ which does not currently offer MS^3^ quantitation. To circumvent this, phosphorylation and ubiquitination PTMs were searched using the Mascot database, and this information was used to manually extract MS^3^ intensities from raw files using the R package mzR.^74^ We manually validated to ensure quality spectral matches, which yielded 13 ubiquitinated lysines and 67 phosphorylated serines or threonines—each with high-confidence spectra across the entire time course. Using scan number and m/z values of MS^2^ from the Mascot results, we extracted intensity values from the matched MS^3^ scans. The Mascot and MaxQuant datasets were joined, and we normalized changes in peptide-level PTMs to underlying changes in total protein abundance.

We saw striking evidence of PTM changes following heat shock. Consistent with the findings of Kanshin et al.,^75^ we saw evidence for dynamic changes in protein phosphorylation levels (Figure 6A). Proteins with at least 1.5-fold increased phosphorylation were enriched for endocytosis (*p* < 3 × 10^−3^), cellular important (*p* < 8 × 10^−3^), and noteably, response to stress (*p* < 8 × 10^−3^). These latter proteins included an enzyme (Tps2p) and regulatory submit (Tps3p) of trehalose biosynthetic complex, both of which are known to be regulated by phosphorylation during stress.^76^ We also observed several proteins with increased or decreased lysine monoubiquitination (Figure 6B). While polyubiquitination has been recognized as a canonical signal for proteasomal degredation, monoubiquitination has recently emerged as another signal for proteasomal degradation.^77^ Intriguingly—and consistent with monubiquitination playing a role as a regulator of protein degradation during heat stress—we observed a strong inverse relationship (*r*^*2*^ = 0.82) between total protein abundance at 90 minutes post-heat shock and fold-change in lysine monoubiquitation at 15 minutes post-heat shock (Figure 6D). Notably, there was no correlation between protein abundance changes and protein phosphorylation changes (*R*^*2*^ = 0.03; Figure 6C), suggesting that the monubiquitation trend is likely biologically meaningful. Overall, our TMT-MS^3^ workflow is precise enough to delineate protein-level and PTM-level changes in biological samples.

**Figure 6.**
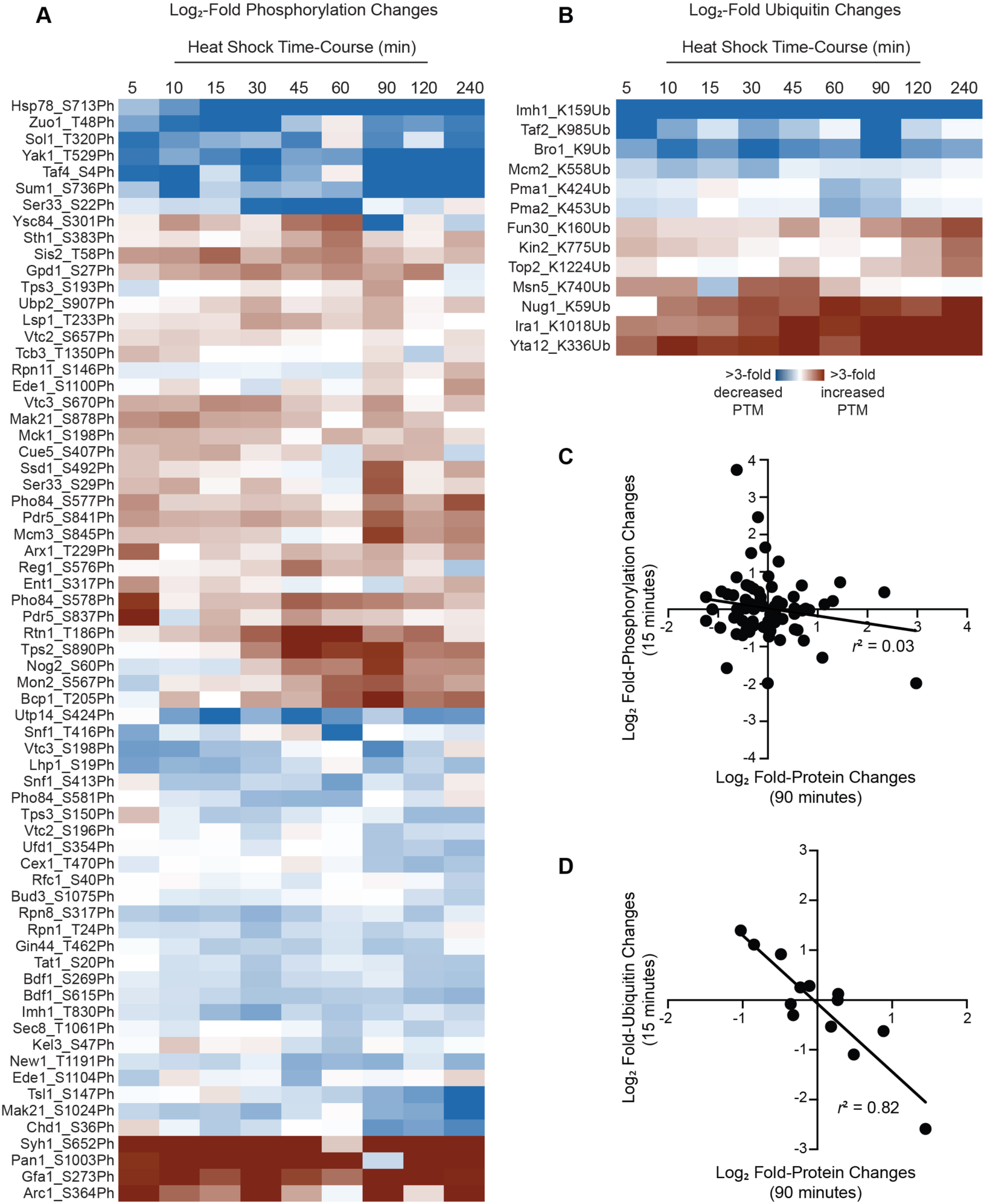
TMT-MS^3^ reveals dynamic changes in the abundance of protein PTMs during heat shock. Modified peptides were identified by searches of MS data using Mascot Distiller and were manually validated as described in the Methods. PTM abundance was normalized to overall abundance changes of the corresponding protein. The heat maps depict heat shock-dependent increases (red) or decreases (blue) in **A)** phosphorylation or **B)** ubiquitination (Note: Pma1_K424Ub and Pma2_K453Ub share the same spectrum). Plot of the correlation between fold-changes in total protein levels and fold changes in **A)** phosphorylation or **B)** ubiquitination.

## Conclusions

In this study, we present a TMT-MS^3^ proteomics workflow that yields precise and accurate measurements of peptide and protein abundance, using the well-studied yeast heat shock response as a test case. At the protein abundance level, our Multinotch MS^3^ analysis method for TMT proteomics was robust to detect differential expression, even in the case of a single replicate. Proteins with significant induction or repression largely recapitulated expectations, with HSP chaperones amongst the most strongly induced proteins, and proteins related to cell growth and protein synthesis among the most strongly repressed. Our method also performed particular well in the quantification of low abundance peptides. Compared to SILAC and label-free methods, the TMT Multinotch MS^3^ workflow affords a significant increase in multiplexing capacity and technical precision for peptide-level quantitative measurements, which is necessary for designing high-power experiments to study temporal dynamics of changing PTMs in a variety of biological contexts. In fact, our analysis of PTMs suggests an additional layer of regulation that has been largely understudied in the context of the yeast heat shock response. While PTMs are known to play key regulatory roles during the adaptation to stress, to date there has been only a single study examining PTM changes during the yeast heat shock response (in this case phosphorylation).^75^ In this study, we identified both phospho- and ubiquitin-modified peptides whose abundance changed during heat shock even following normalization to abundance changes of unmodified peptides for the same protein. We hypothesize that several of the other dynamic PTMs may be important and understudied components of heat shock adaptation. That these were identified without specific enrichment steps suggests that MultiNotch MS^3^ datasets can be mined for PTMs that change in abundance during environmental perturbation.

## Acknowledgements

We thank Nilda Burgos and Reiofeli Salas for the use and assistance with the bead mill, and Shannon Servoss and Neda Mahmoudi for the use and assistance with the lyophilizer. This work was supported by a grant from the National Science Foundation (IOS-1656602) to JAL, by a grant from the National Institutes of Health (GM081766) to WPW, and by the Arkansas Biosciences Institute (Arkansas Settlement Proceeds Act of 2000). REH was partially supported by the University of Arkansas Cell and Molecular Biology Graduate Program. AJT acknowledges support from the National Institutes of Health (R01GM118760, P20GM121293, UL1TR000039, P20GM103625, S10OD018445 and P20GM103429).

---

**Figure S1:**
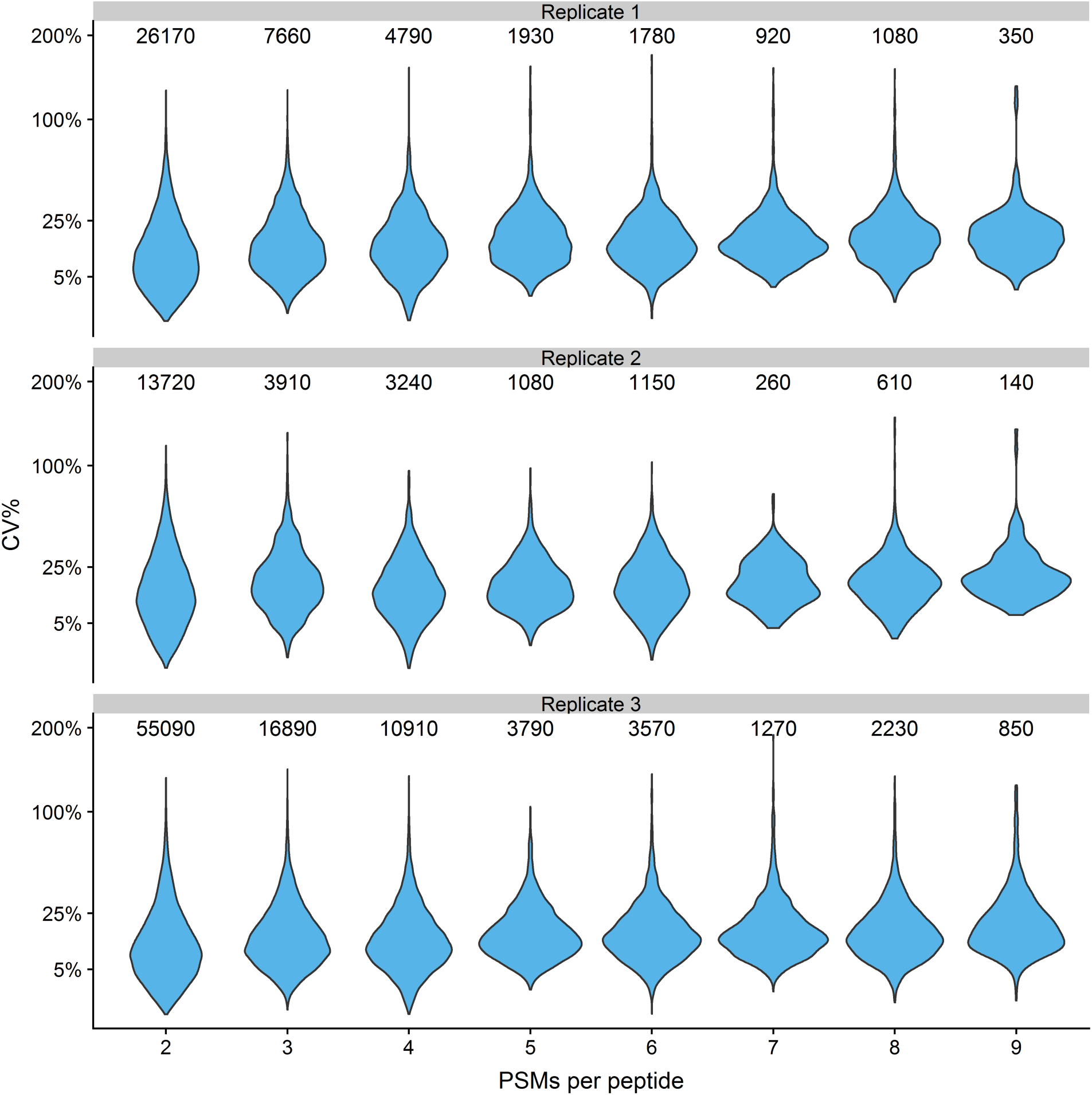
Highly precise, peptide-level measurements for three independent TMT 10-plex experiments. Reporter ion intensity data from three independent TMT 10-plex heat shock time-course experiments were used to calculate coefficient of variance as a function of PSM number.

**Figure S2:**
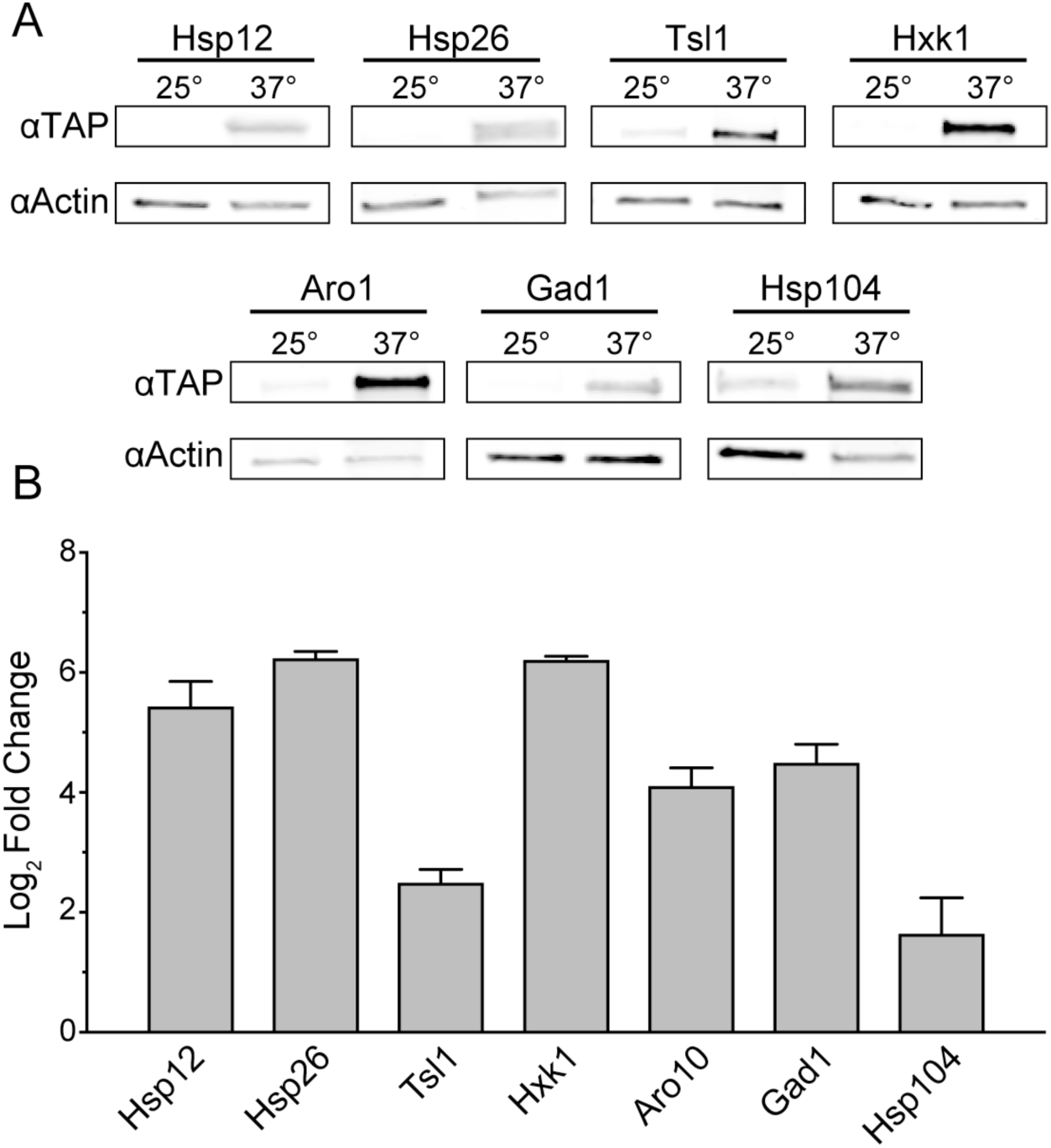
TMT-MS^3^ is an accurate detection method for changes in protein abundance. The relative abundance for seven significantly-induced proteins at 60 minutes post heat-shock was validated via quantitative western blotting. Anti-actin and anti-TAP antibodies were used to detect, simultaneously in the same blot, the Act1p loading control and the TAP-tagged proteins of interest. **A)** Representative western blots. **B)** Quantitation of relative log_2_ fold changes before and after heat shock for each TAP-tagged protein following normalization to actin. Data represent the mean and standard deviation of three biological replicates. Raw images can be found in File S5, and densitometric data can be found in File S6.

**Figure S3:**
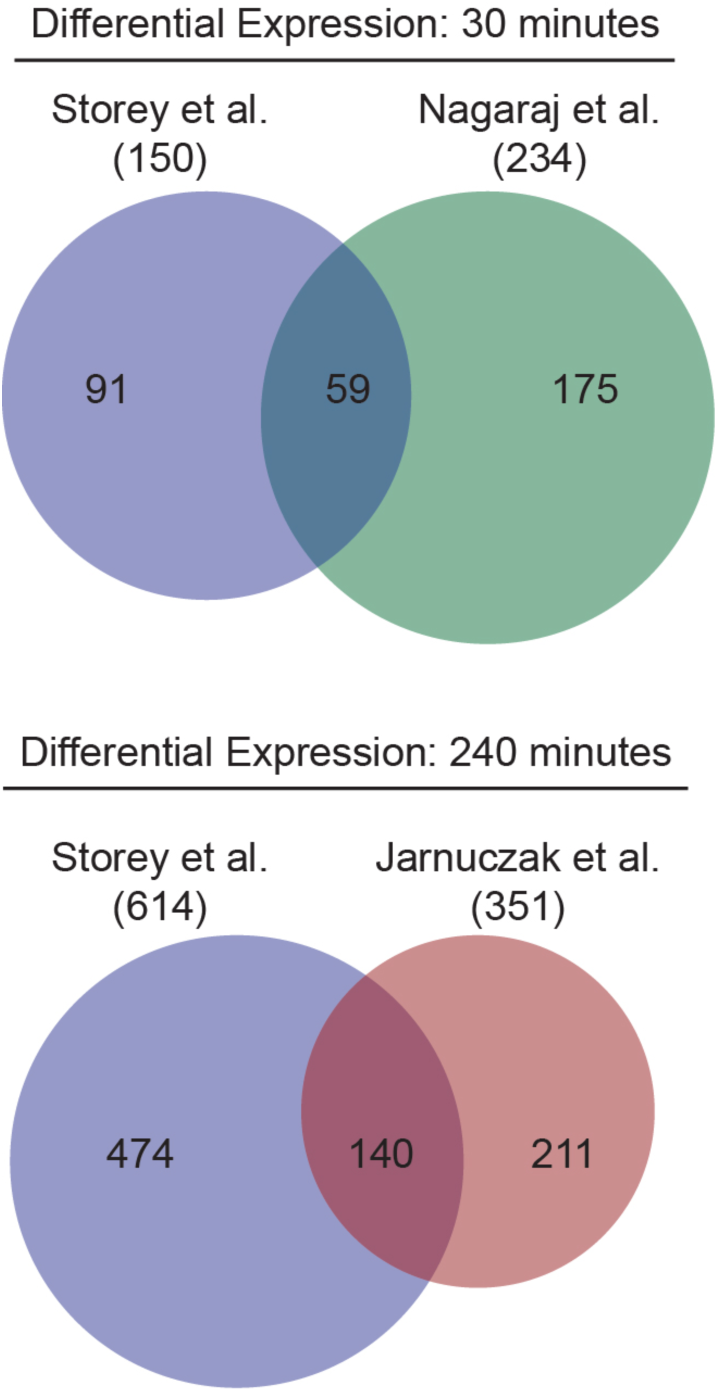
Comparison of differentially abundant proteins from from this TMT-MS^3^ study to those reported by studies which used SILAC and label-free approaches. We compared significantly differentially expressed proteins from this TMT-MS^3^ experiment (Story et al.) to a set of SILAC (Nagaraj et al.^44^) and label-free (Jarnuczak et al.^45^) experiments. Because Nagaraj et al. used an FDR < 0.02, we maintained that threshold for each of these comparisons.

**Figure S4:**
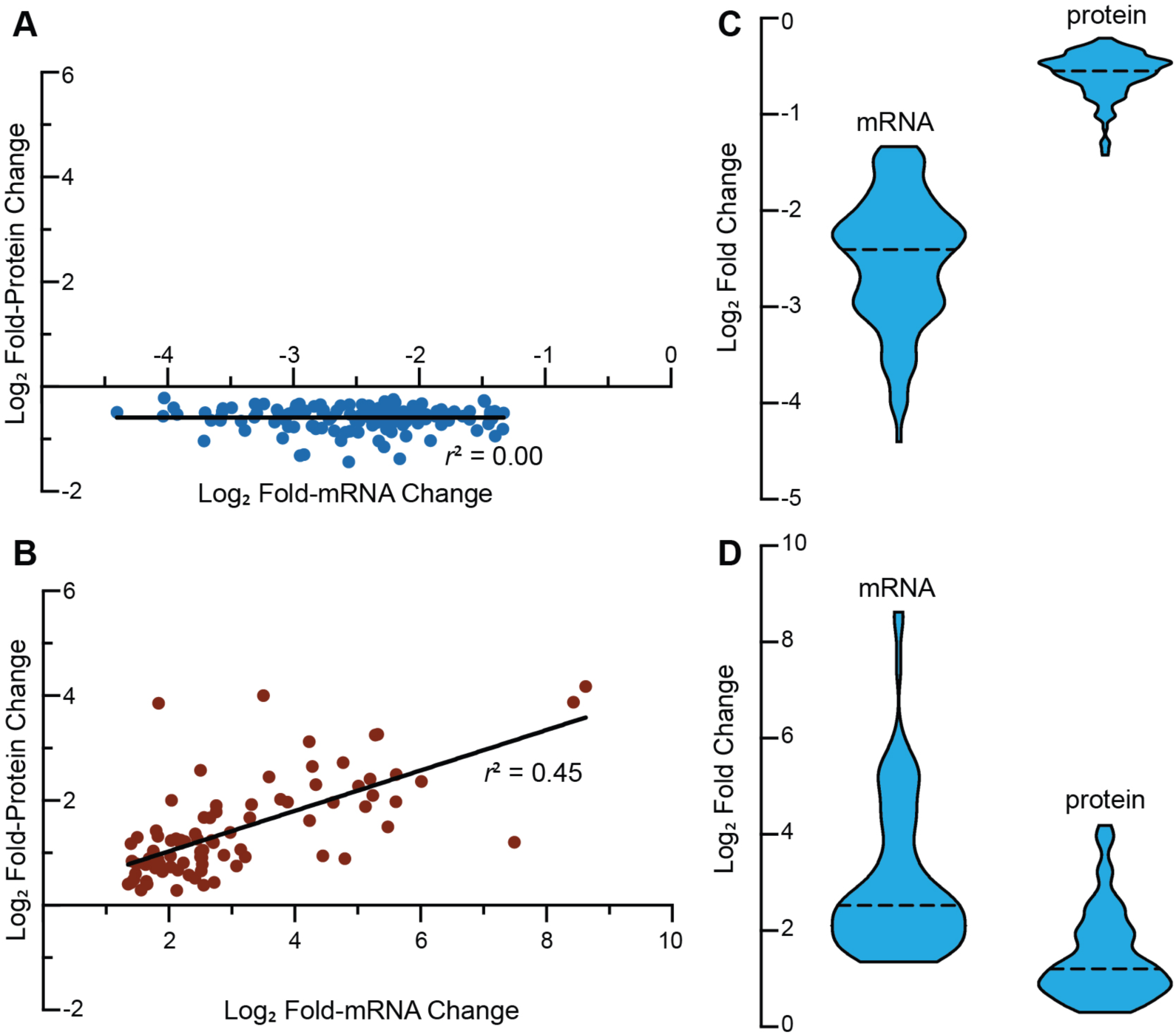
Evidence for buffering of protein repression. Correlation between pairs of proteins and mRNAs from Eng et al.^40^ that were both significantly (FDR < 0.05) **A)** repressed **B)** or induced following heat shock (at 60 minutes for proteins and 15 minutes for mRNAs). The violin plots depict the density of expression values for the **C)** repressed **D)** or induced protein-mRNA pairs.

## Supporting information

**File S1. R scripts used for normalization and analysis.**

**File S2. Limma output for analysis of the heat shock proteome.**

**File S3. Functional enrichments for protein expression clusters.**

**File S4. Regulatory associations for protein expression clusters.**

**File S5. Raw western blot images.**

**File S6. Raw western blot densitometric data.**

## References

(1) Gasch, A. P.; Spellman, P. T.; Kao, C. M.; Carmel-Harel, O.; Eisen, M. B.; Storz, G.; Botstein, D.; Brown, P. O., Genomic expression programs in the response of yeast cells to environmental changes. Molecular biology of the cell 2000, 11 (12), 4241–4257.

(2) Causton, H. C.; Ren, B.; Koh, S. S.; Harbison, C. T.; Kanin, E.; Jennings, E. G.; Lee, T. I.; True, H. L.; Lander, E. S.; Young, R. A., Remodeling of yeast genome expression in response to environmental changes. Molecular biology of the cell 2001, 12, 323–337.

(3) Lee, M. V.; Topper, S. E.; Hubler, S. L.; Hose, J.; Wenger, C. D.; Coon, J. J.; Gasch, A. P., A dynamic model of proteome changes reveals new roles for transcript alteration in yeast. Molecular systems biology 2011, 7, 514.

(4) Zhou, F.; Lu, Y.; Ficarro, S. B.; Adelmant, G.; Jiang, W.; Luckey, C. J.; Marto, J. A., Genome-scale proteome quantification by DEEP SEQ mass spectrometry. Nat Commun 2013, 4, 2171.

(5) Shalon, D.; Smith, S. J.; Brown, P. O., A DNA microarray system for analyzing complex DNA samples using two-color fluorescent probe hybridization. Genome Res 1996, 6 (7), 639–645.

(6) Bentley, D. R.; Balasubramanian, S.; Swerdlow, H. P.; Smith, G. P.; Milton, J.; Brown, C. G.; Hall, K. P.; Evers, D. J.; Barnes, C. L.; Bignell, H. R., et al., Accurate whole human genome sequencing using reversible terminator chemistry. Nature 2008, 456 (7218), 53–59.

(7) Gygi, S. P.; Rochon, Y.; Franza, B. R.; Aebersold, R., Correlation between protein and mRNA abundance in yeast. Molecular and cellular biology 1999, 19 (3), 1720–1730.

(8) Griffin, T. J.; Gygi, S. P.; Ideker, T.; Rist, B.; Eng, J.; Hood, L.; Aebersold, R., Complementary profiling of gene expression at the transcriptome and proteome levels in *Saccharomyces cerevisiae*. Molecular & cellular proteomics: MCP 2002, 1 (4), 323–333.

(9) Nie, L.; Wu, G.; Zhang, W., Correlation between mRNA and protein abundance in *Desulfovibrio vulgaris*: a multiple regression to identify sources of variations. Biochemical and biophysical research communications 2006, 339 (2), 603–610.

(10) Washburn, M. P.; Koller, A.; Oshiro, G.; Ulaszek, R. R.; Plouffe, D.; Deciu, C.; Winzeler, E.; Yates, J. R., 3rd, Protein pathway and complex clustering of correlated mRNA and protein expression analyses in *Saccharomyces cerevisiae*. Proc Natl Acad Sci U S A 2003, 100 (6), 3107–3112.

(11) Csardi, G.; Franks, A.; Choi, D. S.; Airoldi, E. M.; Drummond, D. A., Accounting for experimental noise reveals that mRNA levels, amplified by post-transcriptional processes, largely determine steady-state protein levels in yeast. PLoS genetics 2015, 11 (5), e1005206.

(12) Schwanhausser, B.; Busse, D.; Li, N.; Dittmar, G.; Schuchhardt, J.; Wolf, J.; Chen, W.; Selbach, M., Global quantification of mammalian gene expression control. Nature 2011, 473 (7347), 337–342.

(13) Brar, G. A.; Yassour, M.; Friedman, N.; Regev, A.; Ingolia, N. T.; Weissman, J. S., High-resolution view of the yeast meiotic program revealed by ribosome profiling. Science 2012, 335 (6068), 552–557.

(14) Halbeisen, R. E.; Gerber, A. P., Stress-dependent coordination of transcriptome and translatome in yeast. PLoS Biol 2009, 7 (5), e1000105.

(15) Lackner, D. H.; Schmidt, M. W.; Wu, S.; Wolf, D. A.; Bahler, J., Regulation of transcriptome, translation, and proteome in response to environmental stress in fission yeast. Genome biology 2012, 13 (4), R25.

(16) Medicherla, B.; Goldberg, A. L., Heat shock and oxygen radicals stimulate ubiquitin-dependent degradation mainly of newly synthesized proteins. The Journal of cell biology 2008, 182 (4), 663–673.

(17) Martin-Perez, M.; Villen, J., Determinants and Regulation of Protein Turnover in Yeast. Cell Syst 2017, 5 (3), 283–294 e5.

(18) Brewster, J. L.; de Valoir, T.; Dwyer, N. D.; Winter, E.; Gustin, M. C., An osmosensing signal transduction pathway in yeast. Science 1993, 259 (5102), 1760–1763.

(19) Proft, M.; Pascual-Ahuir, A.; de Nadal, E.; Arino, J.; Serrano, R.; Posas, F., Regulation of the Sko1 transcriptional repressor by the Hog1 MAP kinase in response to osmotic stress. The EMBO journal 2001, 20 (5), 1123–1133.

(20) Han, S. J.; Choi, K. Y.; Brey, P. T.; Lee, W. J., Molecular cloning and characterization of a Drosophila p38 mitogen-activated protein kinase. The Journal of biological chemistry 1998, 273 (1), 369–374.

(21) Rouse, J.; Cohen, P.; Trigon, S.; Morange, M.; Alonso-Llamazares, A.; Zamanillo, D.; Hunt, T.; Nebreda, A. R., A novel kinase cascade triggered by stress and heat shock that stimulates MAPKAP kinase-2 and phosphorylation of the small heat shock proteins. Cell 1994, 78 (6), 1027–1037.

(22) Shivaswamy, S.; Iyer, V. R., Stress-dependent dynamics of global chromatin remodeling in yeast: dual role for SWI/SNF in the heat shock stress response. Molecular and cellular biology 2008, 28 (7), 2221–2234.

(23) Wang, F.; Nguyen, M.; Qin, F. X.; Tong, Q., SIRT2 deacetylates FOXO3a in response to oxidative stress and caloric restriction. Aging cell 2007, 6 (4), 505–514.

(24) Xie, Q.; Hao, Y.; Tao, L.; Peng, S.; Rao, C.; Chen, H.; You, H.; Dong, M. Q.; Yuan, Z., Lysine methylation of FOXO3 regulates oxidative stress-induced neuronal cell death. EMBO Rep 2012, 13 (4), 371–377.

(25) Westerheide, S. D.; Anckar, J.; Stevens, S. M., Jr.; Sistonen, L.; Morimoto, R. I., Stress-inducible regulation of heat shock factor 1 by the deacetylase SIRT1. Science 2009, 323 (5917), 1063–1066.

(26) Magraner-Pardo, L.; Pelechano, V.; Coloma, M. D.; Tordera, V., Dynamic remodeling of histone modifications in response to osmotic stress in *Saccharomyces cerevisiae*. BMC genomics 2014, 15, 247.

(27) Wang, X.; Yen, J.; Kaiser, P.; Huang, L., Regulation of the 26S proteasome complex during oxidative stress. Science signaling 2010, 3 (151), ra88.

(28) Miller, M. J.; Scalf, M.; Rytz, T. C.; Hubler, S. L.; Smith, L. M.; Vierstra, R. D., Quantitative proteomics reveals factors regulating RNA biology as dynamic targets of stress-induced SUMOylation in *Arabidopsis*. Molecular & cellular proteomics: MCP 2013, 12 (2), 449–463.

(29) Golebiowski, F.; Matic, I.; Tatham, M. H.; Cole, C.; Yin, Y.; Nakamura, A.; Cox, J.; Barton, G. J.; Mann, M.; Hay, R. T., System-wide changes to SUMO modifications in response to heat shock. Science signaling 2009, 2 (72), ra24.

(30) Lindquist, S.; Craig, E. A., The heat-shock proteins. Annu Rev Genet 1988, 22, 631–677.

(31) Verghese, J.; Abrams, J.; Wang, Y.; Morano, K. A., Biology of the heat shock response and protein chaperones: budding yeast (*Saccharomyces cerevisiae*) as a model system. Microbiol Mol Biol Rev 2012, 76 (2), 115–158.

(32) Swan, T. M.; Watson, K., Membrane fatty acid composition and membrane fluidity as parameters of stress tolerance in yeast. Can J Microbiol 1997, 43 (1), 70–7.

(33) Mishra, P.; Prasad, R., Alterations in fatty acyl composition can selectively affect amino acid transport in *Saccharomyces cerevisiae*. Biochem Int 1987, 15 (3), 499–508.

(34) Coote, P. J.; Cole, M. B.; Jones, M. V., Induction of increased thermotolerance in *Saccharomyces cerevisiae* may be triggered by a mechanism involving intracellular pH. Journal of general microbiology 1991, 137 (7), 1701–8.

(35) Panaretou, B.; Piper, P. W., The plasma membrane of yeast acquires a novel heat-shock protein (hsp30) and displays a decline in proton-pumping ATPase levels in response to both heat shock and the entry to stationary phase. Eur J Biochem 1992, 206 (3), 635–40.

(36) Davidson, J. F.; Schiestl, R. H., Mitochondrial respiratory electron carriers are involved in oxidative stress during heat stress in *Saccharomyces cerevisiae*. Molecular and cellular biology 2001, 21 (24), 8483–9.

(37) Glover, J. R.; Lindquist, S., Hsp104, Hsp70, and Hsp40: a novel chaperone system that rescues previously aggregated proteins. Cell 1998, 94 (1), 73–82.

(38) Jakob, U.; Gaestel, M.; Engel, K.; Buchner, J., Small heat shock proteins are molecular chaperones. The Journal of biological chemistry 1993, 268 (3), 1517–20.

(39) Singer, M. A.; Lindquist, S., Multiple effects of trehalose on protein folding in vitro and in vivo. Molecular cell 1998, 1 (5), 639–48.

(40) Eng, K. H.; Kvitek, D. J.; Keles, S.; Gasch, A. P., Transient genotype-by-environment interactions following environmental shock provide a source of expression variation for essential genes. Genetics 2010, 184, 587–593.

(41) Wohlbach, D. J.; Rovinskiy, N.; Lewis, J. A.; Sardi, M.; Schackwitz, W. S.; Martin, J. A.; Deshpande, S.; Daum, C. G.; Lipzen, A.; Sato, T. K., et al., Comparative genomics of *Saccharomyces cerevisiae* natural isolates for bioenergy production. Genome biology and evolution 2014, 6, 2557–2566.

(42) Yoon, O. K.; Brem, R. B., Noncanonical transcript forms in yeast and their regulation during environmental stress. RNA 2010, 16, 1256–1267.

(43) Yassour, M.; Pfiffner, J.; Levin, J. Z.; Adiconis, X.; Gnirke, A.; Nusbaum, C.; Thompson, D. A.; Friedman, N.; Regev, A., Strand-specific RNA sequencing reveals extensive regulated long antisense transcripts that are conserved across yeast species. Genome biology 2010, 11, R87.

(44) Nagaraj, N.; Kulak, N. A.; Cox, J.; Neuhauser, N.; Mayr, K.; Hoerning, O.; Vorm, O.; Mann, M., System-wide perturbation analysis with nearly complete coverage of the yeast proteome by single-shot ultra HPLC runs on a bench top Orbitrap. Molecular & cellular proteomics: MCP 2012, 11, M111 013722.

(45) Jarnuczak, A. F.; Albornoz, M. G.; Eyers, C. E.; Grant, C. M.; Hubbard, S. J., A quantitative and temporal map of proteostasis during heat shock in *Saccharomyces cerevisiae*. Mol Omics 2018, 14 (1), 37–52.

(46) Ting, L.; Rad, R.; Gygi, S. P.; Haas, W., MS3 eliminates ratio distortion in isobaric multiplexed quantitative proteomics. Nature methods 2011, 8 (11), 937–940.

(47) McAlister, G. C.; Nusinow, D. P.; Jedrychowski, M. P.; Wuhr, M.; Huttlin, E. L.; Erickson, B. K.; Rad, R.; Haas, W.; Gygi, S. P., MultiNotch MS3 enables accurate, sensitive, and multiplexed detection of differential expression across cancer cell line proteomes. Anal Chem 2014, 86 (14), 7150–7158.

(48) Sun, S.; Zhou, J. Y.; Yang, W.; Zhang, H., Inhibition of protein carbamylation in urea solution using ammonium-containing buffers. Anal Biochem 2014, 446, 76–81.

(49) Wessel, D.; Flugge, U. I., A method for the quantitative recovery of protein in dilute solution in the presence of detergents and lipids. Anal Biochem 1984, 138 (1), 141–143.

(50) Cox, J.; Mann, M., MaxQuant enables high peptide identification rates, individualized p.p.b.-range mass accuracies and proteome-wide protein quantification. Nature biotechnology 2008, 26 (12), 1367–1372.

(51) T., U. C., UniProt: the universal protein knowledgebase. Nucleic acids research 2018, 46 (5), 2699.

(52) Searle, B. C., Scaffold: a bioinformatic tool for validating MS/MS-based proteomic studies. Proteomics 2010, 10 (6), 1265–1269.

(53) Perez-Riverol, Y.; Csordas, A.; Bai, J.; Bernal-Llinares, M.; Hewapathirana, S.; Kundu,. J.; Inuganti, A.; Griss, J.; Mayer, G.; Eisenacher, M., et al., The PRIDE database and related tools and resources in 2019: improving support for quantification data. Nucleic acids research 2019, 47 (D1), D442-D450.

(54) Ritchie, M. E.; Phipson, B.; Wu, D.; Hu, Y.; Law, C. W.; Shi, W.; Smyth, G. K., limma powers differential expression analyses for RNA-sequencing and microarray studies. Nucleic acids research 2015, 43 (7), e47.

(55) Eisen, M. B.; Spellman, P. T.; Brown, P. O.; Botstein, D., Cluster analysis and display of genome-wide expression patterns. Proc Natl Acad Sci U S A 1998, 95, 14863–14868.

(56) Boyle, E. I.; Weng, S.; Gollub, J.; Jin, H.; Botstein, D.; Cherry, J. M.; Sherlock, G., GO::TermFinder--open source software for accessing Gene Ontology information and finding significantly enriched Gene Ontology terms associated with a list of genes. Bioinformatics 2004, 20 (18), 3710–3715.

(57) Teixeira, M. C.; Monteiro, P. T.; Palma, M.; Costa, C.; Godinho, C. P.; Pais, P.; Cavalheiro, M.; Antunes, M.; Lemos, A.; Pedreira, T., et al., YEASTRACT: an upgraded database for the analysis of transcription regulatory networks in Saccharomyces cerevisiae. Nucleic acids research 2018, 46 (D1), D348-D353.

(58) von der Haar, T., Optimized protein extraction for quantitative proteomics of yeasts. PloS one 2007, 2 (10), e1078.

(59) Schneider, C. A.; Rasband, W. S.; Eliceiri, K. W., NIH Image to ImageJ: 25 years of image analysis. Nature methods 2012, 9 (7), 671–675.

(60) Pichler, P.; Kocher, T.; Holzmann, J.; Mazanek, M.; Taus, T.; Ammerer, G.; Mechtler, K., Peptide labeling with isobaric tags yields higher identification rates using iTRAQ 4-plex compared to TMT 6-plex and iTRAQ 8-plex on LTQ Orbitrap. Anal Chem 2010, 82 (15), 6549–6558.

(61) Pottiez, G.; Wiederin, J.; Fox, H. S.; Ciborowski, P., Comparison of 4-plex to 8-plex iTRAQ quantitative measurements of proteins in human plasma samples. Journal of proteome research 2012, 11 (7), 3774–3781.

(62) Corthals, G. L.; Wasinger, V. C.; Hochstrasser, D. F.; Sanchez, J. C., The dynamic range of protein expression: a challenge for proteomic research. Electrophoresis 2000, 21 (6), 1104–1115.

(63) Chen, E. I.; Hewel, J.; Felding-Habermann, B.; Yates, J. R., 3rd, Large scale protein profiling by combination of protein fractionation and multidimensional protein identification technology (MudPIT). Molecular & cellular proteomics: MCP 2006, 5 (1), 53–56.

(64) Wang, Y.; Yang, F.; Gritsenko, M. A.; Wang, Y.; Clauss, T.; Liu, T.; Shen, Y.; Monroe, M. E.; Lopez-Ferrer, D.; Reno, T., et al., Reversed-phase chromatography with multiple fraction concatenation strategy for proteome profiling of human MCF10A cells. Proteomics 2011, 11 (10), 2019–2026.

(65) Mertins, P.; Qiao, J. W.; Patel, J.; Udeshi, N. D.; Clauser, K. R.; Mani, D. R.; Burgess, M. W.; Gillette, M. A.; Jaffe, J. D.; Carr, S. A., Integrated proteomic analysis of post-translational modifications by serial enrichment. Nature methods 2013, 10 (7), 634–637.

(66) Miller, S. B.; Mogk, A.; Bukau, B., Spatially organized aggregation of misfolded proteins as cellular stress defense strategy. J Mol Biol 2015, 427 (7), 1564–1574.

(67) Malinovska, L.; Kroschwald, S.; Munder, M. C.; Richter, D.; Alberti, S., Molecular chaperones and stress-inducible protein-sorting factors coordinate the spatiotemporal distribution of protein aggregates. Molecular biology of the cell 2012, 23 (16), 3041–3056.

(68) Sorger, P. K.; Pelham, H. R., Purification and characterization of a heat-shock element binding protein from yeast. The EMBO journal 1987, 6 (10), 3035–3041.

(69) Martinez-Pastor, M. T.; Marchler, G.; Schuller, C.; Marchler-Bauer, A.; Ruis, H.; Estruch, F., The *Saccharomyces cerevisiae* zinc finger proteins Msn2p and Msn4p are required for transcriptional induction through the stress response element (STRE). The EMBO journal 1996, 15 (9), 2227–2235.

(70) Belle, A.; Tanay, A.; Bitincka, L.; Shamir, R.; O’Shea, E. K., Quantification of protein half-lives in the budding yeast proteome. Proc Natl Acad Sci U S A 2006, 103 (35), 13004–13009.

(71) Liu, Y.; Beyer, A.; Aebersold, R., On the Dependency of Cellular Protein Levels on mRNA Abundance. Cell 2016, 165 (3), 535–550.

(72) Ho, Y. H.; Shishkova, E.; Hose, J.; Coon, J. J.; Gasch, A. P., Decoupling Yeast Cell Division and Stress Defense Implicates mRNA Repression in Translational Reallocation during Stress. Curr Biol 2018, 28 (16), 2673–2680 e4.

(73) Byrum, S. D.; Raman, A.; Taverna, S. D.; Tackett, A. J., ChAP-MS: a method for identification of proteins and histone posttranslational modifications at a single genomic locus. Cell reports 2012, 2 (1), 198–205.

(74) Chambers, M. C.; Maclean, B.; Burke, R.; Amodei, D.; Ruderman, D. L.; Neumann, S.; Gatto, L.; Fischer, B.; Pratt, B.; Egertson, J., et al., A cross-platform toolkit for mass spectrometry and proteomics. Nature biotechnology 2012, 30 (10), 918–920.

(75) Kanshin, E.; Kubiniok, P.; Thattikota, Y.; D’Amours, D.; Thibault, P., Phosphoproteome dynamics of *Saccharomyces cerevisiae* under heat shock and cold stress. Molecular systems biology 2015, 11 (6), 813.

(76) Trevisol, E. T.; Panek, A. D.; de Mesquita, J. F.; Eleutherio, E. C., Regulation of the yeast trehalose-synthase complex by cyclic AMP-dependent phosphorylation. Biochimica et biophysica acta 2014, 1840 (6), 1646–1650.

(77) Braten, O.; Livneh, I.; Ziv, T.; Admon, A.; Kehat, I.; Caspi, L. H.; Gonen, H.; Bercovich, B.; Godzik, A.; Jahandideh, S., et al., Numerous proteins with unique characteristics are degraded by the 26S proteasome following monoubiquitination. Proc Natl Acad Sci U S A 2016, 113 (32), 4639–4647.

