## Supplementary material for "Accurate and Sensitive Quantitation of the Dynamic Heat Shock Proteome using Tandem Mass Tags": File S1: Extract_intensity.R

```
library(tidyverse)
library(magrittr)
library(mzR)

#Goal: Extract reporter ion intensities for Ubiquitinated spectra.

df <- read_tsv("20190306/Spectrum Report for Lewis_mods.xls", skip = 40)
names(df) <- gsub(" ", "_", names(df))

df1 <- df %>%
  filter(grepl("Phospho|GG", Variable_modifications_identified_by_spectrum)) %>%
  select(-c(Experiment_name, Biological_sample_category, Percentage_of_total_spectra,
            Percentage_sequence_coverage, Manual_validation, Assigned)) %>%
  mutate(Raw_file = str_extract(Spectrum_name, "^[^\\.]*")) %>%
  mutate(Raw_file = paste0(Raw_file, ".mzML")) %>%
  separate(Spectrum_name, into = c("Raw", "Scan_number", "Spectrum2", "Charge"),
           remove = FALSE,
           sep = "\\.") %>%
  select(-c(Raw, Spectrum2, Charge)) %>%
  mutate(MS2_scan_number = as.integer(Scan_number))

#Selects Raw file and MS2 scan
extract_info <- df1 %>% select(Raw_file, MS2_scan = MS2_scan_number)

#mzML files with data to be extracted
mzml_files <- unique(extract_info$Raw_file)

#For saving output files
output_names <- gsub("\\.mzML", "", mzml_files)

#Extract intensity function
extract_intensity <- function(df2, mz_value){
  a <- filter(df2, between(V1, mz_value - 0.003, mz_value + 0.003))
  if(length(a$V1) > 0){
    max(a$V2)}
  else{0}
}

i=1

#For each raw file, filters extract_info for MS2 scans.  Opens corresponding mzML file.
#Finds MS3 scans with precursor scan number that matches MS2 scan.  Extracts full scan data
#For each report ion, finds max intensity value within 0.003 mz of the TMT reporter ion mz.
#Writes to "Extracted_intensity_for" + file name.

output_vector <- list()

for(i in seq_along(mzml_files)){
  
  file_scans <- filter(extract_info, Raw_file == mzml_files[[i]]) %>%
    distinct()
  mzml <- openMSfile(str_c("../raw_files/mzML_files/", mzml_files[[i]]))
  mzml_header <- header(mzml) %>%
    mutate(Raw_file = mzml_files[[i]]) %>%
    filter(msLevel == 3,
           precursorScanNum %in% file_scans$MS2_scan) %>%
    select(Raw_file, MS2_scan = precursorScanNum, seqNum) %>%
    mutate(peak_data = map(seqNum, ~peaks(mzml, scans = .x) %>%
                             as.data.frame() %>%
                             filter(., between(V1, 125, 133))))
  
  a2 <- mzml_header %>%
    mutate(TMT_126  = map_dbl(peak_data, ~extract_intensity(.x, 126.127726)),
           TMT_127N = map_dbl(peak_data, ~extract_intensity(.x, 127.124761)),
           TMT_127C = map_dbl(peak_data, ~extract_intensity(.x, 127.131081)),
           TMT_128N = map_dbl(peak_data, ~extract_intensity(.x, 128.128116)),
           TMT_128C = map_dbl(peak_data, ~extract_intensity(.x, 128.134436)),
           TMT_129N = map_dbl(peak_data, ~extract_intensity(.x, 129.131471)),
           TMT_129C = map_dbl(peak_data, ~extract_intensity(.x, 129.13779)),
           TMT_130N = map_dbl(peak_data, ~extract_intensity(.x, 130.134825)),
           TMT_130C = map_dbl(peak_data, ~extract_intensity(.x, 130.141145)),
           TMT_131  = map_dbl(peak_data, ~extract_intensity(.x, 131.13818))) %>%
    select(-peak_data)
  
  
  output_vector[[i]] <- a2
  print(paste(output_names[[i]], "done!"))
  
  close(mzml)
  
}

a3 <- do.call(rbind, output_vector)


df4 <- df1 %>%
  rename(MS2_scan = MS2_scan_number) %>%
  left_join(a3)

write_tsv(df4, "20190306/Extracted_intensity.txt")
```
