## Supplementary material for "Accurate and Sensitive Quantitation of the Dynamic Heat Shock Proteome using Tandem Mass Tags": File S1: Figure_2.R

```
library(tidyverse)
library(cowplot)
library(ggsci)


theme_set(theme_cowplot(font_size=12)) # reduce default font size

df1<-read_tsv("raw/Maxquant_txt/evidence.txt", guess_max = 30000)
names(df1) <- gsub(" ", "_", names(df1))


# Figure 1 and 2 Tidy ------------------------------------------------------------

df <- df1 %>%
  filter(is.na(Potential_contaminant),
         is.na(Reverse)) %>%
  mutate(Foldchange = Reporter_intensity_1 / Reporter_intensity_0,
         Foldchange = log2(Foldchange))  %>%
  filter(!is.na(Foldchange),
         !is.infinite(Foldchange)) %>%
  mutate(Foldchange = as.numeric(scale(Foldchange)))

evidence_df_fraction <- df %>% group_by(Peptide_ID) %>% summarize(Resolution=n_distinct(Raw_file))

evidence_df_fraction <-evidence_df_fraction %>% 
  mutate(Resolution = ifelse(Resolution > 5, ">5", Resolution),
         Resolution = factor(Resolution, levels = c(1,2,3,4,5, ">5")))

evidence_df_fraction_num <- evidence_df_fraction %>%
  group_by(Resolution) %>%
  summarize(num=n(),
            percent_total=n()/199.14,
            percent_total=round(percent_total, digit = 1))

Peptide_df<- df %>%
  group_by(Peptide_ID) %>% 
  summarize (Peptide_count =n(),
             Peptide_avg = mean(2^Foldchange,na.rm=TRUE),
             Peptide_sd =  sd(2^Foldchange),
             Peptide_cv = Peptide_sd / Peptide_avg) %>%
  ungroup() %>%
  filter (Peptide_count>1) %>%
  mutate(Count_factor = ifelse(Peptide_count > 6, ">6", Peptide_count),
         Count_factor = factor(Count_factor, levels = c("2", "3", "4", "5", "6", ">6")))

Peptide_count_data <- Peptide_df %>% group_by(Count_factor) %>% summarize(data_count=n(),max_value=max(Peptide_sd))

evidence_df_fraction2 <- df %>% group_by(Raw_file) %>% summarize(Peptide_count = n_distinct(Peptide_ID)) %>%
  mutate(Superfraction = as.integer(gsub(".*SF24\\_", "", Raw_file)))


# Figure 2 Construction ---------------------------------------------------

(gg_fraction <- ggplot(evidence_df_fraction)+
   geom_bar(aes(x=Resolution),fill="#56B4E9", color = "black") +
   geom_text(data=evidence_df_fraction_num,aes(x=Resolution,y=num+1000,label=paste(percent_total,"%",sep="")))+
   #geom_text(data=evidence_df_fraction_num,aes(x=Resolution,y=num+3000,label=num))+
   labs(x="Number of fractions",y="Peptides"))

(gg2 <- evidence_df_fraction2 %>%
    ggplot(aes(x = as.factor(Superfraction), y = Peptide_count)) +
    geom_bar(stat = "identity", fill="#56B4E9", color = "black") +
    labs(x = "Superfraction", y = "Peptides"))

(F1 <- plot_grid(gg_fraction, gg2, nrow = 1, rel_widths = c(.5, 1), labels = c("A", "B")))


save_plot("plots/Figure2.tiff", F1,
          nrow = 1, ncol = 1, base_width = 10, base_height = 3.5,
          compression = "lzw", dpi = 600)
```
