## Supplementary material for "Accurate and Sensitive Quantitation of the Dynamic Heat Shock Proteome using Tandem Mass Tags": File S1: Figure_3.R

```
library(tidyverse)
library(cowplot)
library(ggsci)

theme_set(theme_cowplot(font_size=12)) # reduce default font size

df1<-read_tsv("raw/Maxquant_txt/evidence.txt", guess_max = 30000)
names(df1) <- gsub(" ", "_", names(df1))


# Tidy ------------------------------------------------------------

df <- df1 %>%
  filter(is.na(Potential_contaminant),
         is.na(Reverse)) %>%
  mutate(Foldchange = Reporter_intensity_1 / Reporter_intensity_0,
         Foldchange = log2(Foldchange))  %>%
  filter(!is.na(Foldchange),
         !is.infinite(Foldchange)) %>%
  mutate(Foldchange = as.numeric(scale(Foldchange)))

evidence_df_fraction <- df %>% group_by(Peptide_ID) %>% summarize(Resolution=n_distinct(Raw_file))

evidence_df_fraction <-evidence_df_fraction %>% 
  mutate(Resolution = ifelse(Resolution > 5, ">5", Resolution),
         Resolution = factor(Resolution, levels = c(1,2,3,4,5, ">5")))

evidence_df_fraction_num <- evidence_df_fraction %>%
  group_by(Resolution) %>%
  summarize(num=n(),
            percent_total=n()/199.14,
            percent_total=round(percent_total, digit = 1))

Peptide_df<- df %>%
  group_by(Peptide_ID) %>% 
  summarize (Peptide_count =n(),
             Peptide_avg = mean(2^Foldchange,na.rm=TRUE),
             Peptide_sd =  sd(2^Foldchange),
             Peptide_cv = Peptide_sd / Peptide_avg) %>%
  ungroup() %>%
  filter (Peptide_count>1) %>%
  mutate(Count_factor = ifelse(Peptide_count > 6, ">6", Peptide_count),
         Count_factor = factor(Count_factor, levels = c("2", "3", "4", "5", "6", ">6")))

Peptide_count_data <- Peptide_df %>% group_by(Count_factor) %>% summarize(data_count=n(),max_value=max(Peptide_sd))


df_scatter<-df %>%
  group_by(Peptide_ID) %>%
  mutate (Peptide_count=n(),Peptide_avg=mean(Foldchange,na.rm=TRUE),Peptide_sd=sd(Foldchange)) %>% filter (Peptide_count==2)
df_scatter$Spread_key <- c(1:2)
df_scatter <- df_scatter %>% select (Peptide_ID,Foldchange,Peptide_sd, Spread_key) %>% spread(Spread_key,Foldchange)
quantile(df_scatter$Peptide_sd,c(.95,.98,.99))


a <- lm(`2` ~ `1`, data = filter(df_scatter, Peptide_sd < .9941736))
summary(a)

a$coefficients

a <- lm(`2` ~ `1`, data = df_scatter)
summary(a)


# Figure 3 Construction ---------------------------------------------------
(gg_violin <- Peptide_df %>%
   ggplot() +
   geom_violin(aes(x = Count_factor,y=Peptide_cv*100), fill="#56B4E9") +
   scale_y_continuous(trans="sqrt",
                      breaks = c(5, 25, 100, 200),
                      labels = c("5%", "25%", "100%", "200%")) +
   geom_text(data=Peptide_count_data,
             aes(x=Count_factor, y = 400, label=data_count, vjust = "top")) +
   labs(x="PSMs per peptide",y="CV%"))

(gg_FC <- ggplot(df_scatter,aes(x=`1`,y=`2`))+
    geom_point(alpha = .1)+
    geom_abline(slope = 0.93379, intercept = 0.008618, alpha = .5) +
    labs(x=expression("Log"[2]*" fold change 1"),
         y=expression("Log"[2]*" fold change 2"))
  #annotate("text", x = 5, y = 7.5, label = expression("r"^"2"*" = 0.88"))
)

(F2 <- plot_grid(gg_violin, gg_FC, nrow = 2, labels = c("A", "B"), align = "v"))


# Figure 3 Save -----------------------------------------------------------
save_plot("plots/Figure3.pdf", F2,
          nrow = 2, ncol = 2, base_width = 3, base_height = 3,
          paper = "letter")

save_plot("plots/Figure3.tiff", F2,
          nrow = 2, ncol = 2, base_width = 3, base_height = 3,
          compression = "lzw", dpi = 600)
```
