## Supplementary material for "Accurate and Sensitive Quantitation of the Dynamic Heat Shock Proteome using Tandem Mass Tags": File S1: Fix PTM function.R

```
fix_PTMs <- function(df){
  
  #If histone, subtracts 1 from Peptide start index.
  df$Peptide_start_index <- if_else(grepl("Histone H", df$Protein_name),
                                                              df$Peptide_start_index-1L,
                                                              df$Peptide_start_index)
  #Filters for rows with modifications (excludes TMT)
  a1 <- df %>%
    filter(!is.na(Variable_modifications_identified_by_spectrum))
  a2 <- df %>%
    filter(is.na(Variable_modifications_identified_by_spectrum))
  
  #Replace n-term with placeholder
  a1 <- a1 %>%
    mutate(Variable_modifications_identified_by_spectrum = str_replace_all(Variable_modifications_identified_by_spectrum,
                                                                           "n\\-term", "x1"))
  
  
  a <- str_extract_all(a1$Variable_modifications_identified_by_spectrum,
                       "[[:alpha:]][[:digit:]]+\\: [:alpha:]+")
  
  a_AA <- str_extract_all(a, "[[:alpha:]](?=[[:digit:]]+\\: )")
  
  a_site <- str_extract_all(a, "(?<=[[:alpha:]])[[:digit:]]+(?=\\: )") %>%
    map(as.numeric) %>%
    map2(a1$Peptide_start_index-1, ~(.x + .y)) %>%
    map(as.character)
  
  a_mod <- str_extract_all(a,"(?<=[[:digit:]]{1,3})\\: [[:alpha:]]*")
  
  a4 <- map2(a_AA, a_site, ~str_c(.x, .y)) %>%
    map2(a_mod, ~str_c(.x, .y))
  
  a5 <- a4 %>%
    map_chr(~paste(., collapse = ", "))
  
  Acetyl_sites <- a4 %>%
    map(~grep("Acetyl", .x, value = TRUE)) %>%
    map_chr(~paste(., collapse = ", "))
  
  Oxidation_sites <- a4 %>%
    map(~grep("Oxidation", .x, value = TRUE)) %>%
    map_chr(~paste(., collapse = ", "))
  
  GG_sites <- a4 %>%
    map(~grep("GG", .x, value = TRUE)) %>%
    map_chr(~paste(., collapse = ", "))
  
  Phospho_sites <- a4 %>%
    map(~grep("Phospho", .x, value = TRUE)) %>%
    map_chr(~paste(., collapse = ", "))
  
  a1$correct_PTM <- a5 %>%
    str_replace_all("x1", "n-term") %>%
    str_replace_all("x2", "n-term")
  a1$Variable_modifications_identified_by_spectrum <- str_replace_all(a1$Variable_modifications_identified_by_spectrum,
                                                                      "x1", "n-term")
  a1$Acetyl_sites <- Acetyl_sites %>%
    str_replace_all("x1", "n-term") %>%
    str_replace_all("x2", "n-term")
  
  a1$Oxidation_sites <- Oxidation_sites %>%
    str_replace_all("x1", "n-term") %>%
    str_replace_all("x2", "n-term")
  
  a1$GG_sites <- GG_sites %>%
    str_replace_all("x1", "n-term") %>%
    str_replace_all("x2", "n-term")
  
  a1$Phospho_sites <- Phospho_sites %>%
    str_replace_all("x1", "n-term") %>%
    str_replace_all("x2", "n-term")
  
  
  a2$correct_PTM <- NA_character_
  a2$Acetyl_sites <- NA_character_
  a2$Oxidation_sites <- NA_character_
  a2$GG_sites <- NA_character_
  a2$Phospho_sites <- NA_character_
  
  return(rbind(a1, a2))
  
}
```
