## Supplementary material for "Accurate and Sensitive Quantitation of the Dynamic Heat Shock Proteome using Tandem Mass Tags": File S1: Make summary table for heat shock paper.R

```
library(tidyverse)
library(ggsci)
library(broom)
library(cowplot)

#tables
time_point_table <- read_tsv("bin/Time_point_table.txt")
raw_file_table <- read_tsv("bin/raw_file_table.txt")

df5 <- read_tsv("data/S3_Transform.txt", guess_max = 100000)

#Read in transformed data
PTM_table <- read_tsv("data/tables/PTM_id_table.txt")
time_point_table <- read_tsv("bin/Time_point_table.txt")

#Read in list relating Gene name to Systematic name (For Dr. Lewis)
Gene_name_table <- read_tsv("bin/Yeast_Gene_List.txt") %>%
  select(-`Gene Name`)
names(Gene_name_table) <- c("Systematic_name", "Gene_name")

#Reads Protein_id_table, joins with Systematic name
Prot_table <- read_tsv("data/tables/Protein_id_table.txt") %>%
  mutate(Gene_name = gsub("^.*GN\\=", "", Protein_name) %>% gsub(" PE\\=.*$", "", .)) %>%
  left_join(Gene_name_table)

#This contains all intensity values (1-4) for all spectra used in downstream analysis.  This will be used to relate interesting findings to the raw data.
tidy_df <- df5 %>%
  mutate(Protein_id = as.integer(Protein_id)) %>%
  left_join(Prot_table) %>%
  #Join protein name with PTM site
  mutate(correct_PTM1 = gsub("\\: Acetyl", "Ac", correct_PTM) %>%
           gsub("\\: Oxidation", "Ox", .),
         correct_PTM1 = ifelse(is.na(correct_PTM1), "Unmodified", correct_PTM1)) %>%
  mutate(Protein_accession_numbers = gsub("\\_YEAST", "", Protein_accession_numbers)) %>%
  unite(Prot_PTM, Protein_accession_numbers, correct_PTM1) %>%
  ungroup() %>%
  select(Replicate, Raw_file, Spectrum_id, Protein_id, PTM_id, Peptide_id, Protein, Gene_name, Systematic_name, correct_PTM, Prot_PTM, Acetyl_sites,
         Oxidation_sites, Time, Intensity, Intensity2, Intensity3, Intensity4, Intensity5)


df <- tidy_df %>%
  select(PTM_id, Replicate, Time, Intensity3) %>%
  group_by(Replicate, PTM_id, Time) %>%
  mutate(Intensity3 = 2^Intensity3,
         Count = n()) %>%
  filter(Count > 1) %>%
  summarize(Count = n(),
            Mean = mean(Intensity3),
            SD = sd(Intensity3),
            CV = SD/Mean)

write_tsv(df, "20190306/data/CV_summary_timecourse.txt")

df %>%
  filter(Count < 30) %>%
  ggplot(aes(x = as.factor(Count), y = CV)) + geom_violin()

df1 <- df  %>%
  filter(SD > 0,
         Count < 10)

Count_data <- df1 %>%
  group_by(Count, Replicate) %>%
  summarize(A = n())


(gg_violin <- df1 %>%
    filter(SD > 0) %>%
    ggplot() +
    geom_violin(aes(x = as.factor(Count), y=CV*100), fill="#56B4E9") +
    scale_y_continuous(trans="sqrt",
                       breaks = c(5, 25, 100, 200),
                       labels = c("5%", "25%", "100%", "200%")) +
    geom_text(data = Count_data, aes(label = A, x = as.factor(Count), y = 200)) +
    facet_wrap(~paste("Replicate",Replicate), nrow = 3) +
    labs(x="PSMs per peptide",y="CV%"))
```
