## Supplementary material for "Accurate and Sensitive Quantitation of the Dynamic Heat Shock Proteome using Tandem Mass Tags": File S1: S2_Tidy.R

```
library(tidyverse)
library(ggsci)
source("src/Fix PTM function.R")

#tables
time_point_table <- read_tsv("bin/Time_point_table.txt")
raw_file_table <- read_tsv("bin/raw_file_table.txt")


df <- read_tsv("raw/S1_Mascot_report.txt", guess_max = 50000) %>%
  select(-TMT_131C)
df_mods <- read_tsv("20190306/Extracted_intensity.txt") %>%
  rename(MS2_scan_number = MS2_scan,
         MS3_scan_number = seqNum,
         TMT_131N = TMT_131) %>%
  mutate(`MS/MS_sample_name` = gsub("Mudpit\\_", "", `MS/MS_sample_name`))

df <- rbind(df, df_mods)

dfa <- df %>%
  filter(!grepl("DECOY", Protein_name)) %>%
  fix_PTMs() %>%
  mutate(Gene_name = gsub("^.* GN\\=", "", Protein_name) %>% gsub(" PE\\=.*$", "", .),
         Protein = gsub(" OS\\=.*$", "", Protein_name),
         correct_PTM = str_c(Acetyl_sites, Oxidation_sites, GG_sites, Phospho_sites, sep = ", ") %>%
           gsub("\\, $", "", .) %>%
           gsub("^\\, ", "", .)) %>%
  ungroup()


#SQL-like tables
df1 <- dfa %>%
  select(-c(Database_sources, `Actual_minus_calculated_peptide_mass_(AMU)`, `Actual_minus_calculated_peptide_mass_(PPM)`,
            `Retention_Time_(sec)`, Total_Ion_Current)) %>%
  rowid_to_column("Spectrum_id") %>%
  filter(!is.na(Biological_sample_name)) %>%
  write_tsv("20190306/data/tables/Spectrum_id_table.txt")

Protein_table <- df1 %>%
  distinct(Protein_name, Protein, Protein_accession_numbers, `Protein_molecular_weight_(Da)`) %>%
  rowid_to_column("Protein_id") %>%
  write_tsv("20190306/data/tables/Protein_id_table.txt")

PTM_table <- df1 %>%
  distinct(Peptide_sequence, Fixed_modifications_identified_by_spectrum, Variable_modifications_identified_by_spectrum,
           correct_PTM, Peptide_start_index,
           Peptide_stop_index, Exclusive,
           Acetyl_sites) %>%
  rowid_to_column("PTM_id") %>%
  write_tsv("20190306/data/tables/PTM_id_table.txt")

Peptide_table <- df1 %>%
  distinct(
    Protein_name,
    Protein_accession_numbers,
    Peptide_sequence,
    Fixed_modifications_identified_by_spectrum,
    Variable_modifications_identified_by_spectrum,
    `Calculated_+1H_Peptide_Mass_(AMU)`
  ) %>%
  rowid_to_column("Peptide_id") %>%
  write_tsv("20190306/data/tables/Peptide_id_table.txt")

Tidy_with_ids <- df1 %>%
  left_join(Protein_table) %>%
  left_join(PTM_table) %>%
  left_join(Peptide_table)

df <- Tidy_with_ids %>%
  right_join(raw_file_table, .) %>%
  write_tsv("20190306/data/S2_Tidy.txt")
```
