## Supplementary material for "Accurate and Sensitive Quantitation of the Dynamic Heat Shock Proteome using Tandem Mass Tags": File S1: S3_Transform.R

```
library(tidyverse)
library(ggsci)
library(cowplot)
theme_set(theme_cowplot(font_size=12)) # reduce default font size


#tables
time_point_table <- read_tsv("bin/Time_point_table.txt")
raw_file_table <- read_tsv("bin/raw_file_table.txt")


#Makes data folder if it doesn't already exist
make_folders <- function(x1){
  if(!dir.exists(x1)){
    dir.create(x1)
  }
}

make_folders("20190306/plots")
make_folders("20190306/plots/normalization")

df <- read_tsv("20190306/data/S2_Tidy.txt", guess_max = 100000) %>%
  mutate(Replicate = ifelse(is.na(Replicate), as.integer(gsub("Rep", "", Biological_sample_name)), Replicate),
         Enrichment = ifelse(is.na(Enrichment), "Unenriched", Enrichment),
         Raw_file = gsub("\\.mzML", ".raw", Raw_file))


# Filter ------------------------------------------------------------------

df1 <- df %>%
  #filter(Mascot_Ion_score > 15) %>% #14478 rows lost
  #filter(!(Enrichment == "Enriched" & is.na(Acetyl_sites))) %>% # 21585 rows lost
  gather(Channel, Intensity, TMT_126:TMT_131N) %>%
  left_join(time_point_table) %>%
  mutate(Intensity = ifelse(Intensity > 0, log2(Intensity), Intensity)) %>%
  mutate(Time = factor(Time, levels = time_point_table$Time),
         Channel = factor(Channel, levels = time_point_table$Channel)) %>%
  group_by(Spectrum_id) %>%
  filter(any(Intensity > 0)) %>%
  ungroup() %>%
  filter(Enrichment == "Unenriched")

# Normalization -----------------------------------------------------------


#Transformation 1: Z-score
#Transformation 2: Adjust mixing
#Transformation 3: Scale to average unmodified protein value at time point 0
#Transformation 4: Scale to average unmodified protein value for all time points

#Transformation 1 and 2
df2 <- df1  %>%
  group_by(Spectrum_id) %>%
  filter(all(Intensity > 0)) %>%
  group_by(Spectrum_id) %>%
  mutate(Intensity2 = Intensity -mean(Intensity)) %>% #Center spectrum
  mutate(Acetylated = ifelse(!is.na(Acetyl_sites), "Acetylated", "Unacetylated")) %>% #Will adjust for mixing
  #mutate(Intensity2 = as.numeric(scale(Intensity))) %>% #Scale spectrum
  group_by(Channel, Replicate) %>% #Adjust for mixing, assumes all channels should have equal medians
  mutate(Intensity3 = Intensity2 - median(Intensity2)) %>% 
  #mutate(Intensity3 = Intensity2) %>% #Don't adjust for mixing
  ungroup() %>%
  select(-Acetylated)

#Transformation 3.
Prot_df2 <- df2 %>%
  filter(is.na(correct_PTM)) %>%
  group_by(Protein_id, Time, Replicate) %>%
  summarize(Avg_Intensity3 = mean(Intensity3)) %>%
  ungroup()


#Transformation 4
Prot_df2_time0 <- Prot_df2 %>%
  filter(Time == 0) %>%
  select(-Time)

df3 <- df2 %>%
  left_join(Prot_df2_time0) %>%
  mutate(Avg_Intensity3 = ifelse(is.na(Avg_Intensity3), 0, Avg_Intensity3)) %>%
  ungroup() %>%
  mutate(Intensity4 = Intensity3 - Avg_Intensity3)

df4 <- df3 %>%
  select(-Avg_Intensity3) %>%
  right_join(Prot_df2) %>%
  mutate(Intensity5 = Intensity3 - Avg_Intensity3)


df5 <- df4 %>%
  select(Replicate, matches("\\_id$"), Raw_file, Enrichment, Acetyl_sites, Oxidation_sites, GG_sites, Phospho_sites, correct_PTM,
         Time,  Channel, matches("Intensity"), -Avg_Intensity3) %>%
  ungroup() %>%
  write_tsv("20190306/data/S3_Transform.txt")
```
