## Supplementary material for "Accurate and Sensitive Quantitation of the Dynamic Heat Shock Proteome using Tandem Mass Tags": File S1: S4_Summarize.R

```
library(tidyverse)
library(ggsci)
library(broom)
library(cowplot)

#tables
time_point_table <- read_tsv("bin/Time_point_table.txt")
raw_file_table <- read_tsv("bin/raw_file_table.txt")

df5 <- read_tsv("20190306/data/S3_Transform.txt", guess_max = 100000)

#Read in transformed data
PTM_table <- read_tsv("20190306/data/tables/PTM_id_table.txt")
time_point_table <- read_tsv("bin/Time_point_table.txt")

#Read in list relating Gene name to Systematic name (For Dr. Lewis)
Gene_name_table <- read_tsv("bin/Yeast_Gene_List.txt") %>%
  select(-`Gene Name`)
names(Gene_name_table) <- c("Systematic_name", "Gene_name")

#Reads Protein_id_table, joins with Systematic name, saves to tsv
Prot_table <- read_tsv("20190306/data/tables/Protein_id_table.txt") %>%
  mutate(Gene_name = gsub("^.*GN\\=", "", Protein_name) %>% gsub(" PE\\=.*$", "", .)) %>%
  left_join(Gene_name_table)

#This contains all intensity values (1-4) for all spectra used in downstream analysis.  This will be used to relate interesting findings to the raw data.
tidy_df <- df5 %>%
  mutate(Protein_id = as.integer(Protein_id)) %>%
  left_join(Prot_table) %>%
  #Join protein name with PTM site
  mutate(correct_PTM1 = gsub("\\: Acetyl", "Ac", correct_PTM) %>%
           gsub("\\: Oxidation", "Ox", .),
         correct_PTM1 = ifelse(is.na(correct_PTM1), "Unmodified", correct_PTM1)) %>%
  mutate(Protein_accession_numbers = gsub("\\_YEAST", "", Protein_accession_numbers)) %>%
  unite(Prot_PTM, Protein_accession_numbers, correct_PTM1) %>%
  ungroup() %>%
  select(Replicate, Raw_file, Spectrum_id, Protein_id, PTM_id, Peptide_id, Protein, Gene_name, Systematic_name, correct_PTM, Prot_PTM, Acetyl_sites,
         Oxidation_sites, GG_sites, Phospho_sites, Time, Intensity, Intensity2, Intensity3, Intensity4, Intensity5) %>%
  write_tsv("20190306/data/S6_tidy_all_evidence.txt")

#Normalized protein abundance.  Running this with Intensity5 yields intensity values equal or very close to 0 for all cases (as it should).
Protein_summary_df <- tidy_df %>%
  filter(is.na(correct_PTM)) %>%
  group_by(Protein_id, Time, Replicate, Protein, Systematic_name, Gene_name) %>%
  summarize(Avg_Intensity4 = mean(Intensity4)) %>%
  spread(Time, Avg_Intensity4) %>%
  write_tsv("20190306/data/S6_summarized_protein_abundance.txt")


#Acetyl peptides, normalized by protein expression.  Does a good job removing oxidated peptides.
normalized_acetyl_df <- tidy_df %>%
  filter(!is.na(Acetyl_sites)) %>%
  group_by(Protein, Systematic_name, Prot_PTM, Replicate, Time) %>%
  summarize(Intensity = mean(Intensity5)) %>%
  group_by(Replicate, Time) %>%
  mutate(Intensity = Intensity - median(Intensity)) %>%
  ungroup() %>%
  spread(Time, Intensity) %>%
  write_tsv("20190306/data/S6_normalized_summarized_acetyl_abundance.txt")


#Acetyl peptides, NOT normalized by protein expression.  Does a good job removing oxidated peptides.
acetyl_df <- tidy_df %>%
  filter(!is.na(Acetyl_sites)) %>%
  group_by(Prot_PTM, Replicate) %>%
  filter(sum(Intensity4) != 0) %>%
  group_by(Protein, Systematic_name, Prot_PTM, Replicate, Time) %>%
  summarize(Intensity = mean(Intensity4)) %>%
  group_by(Replicate, Time) %>%
  mutate(Intensity = Intensity - median(Intensity)) %>%
  ungroup() %>%
  spread(Time, Intensity) %>%
  write_tsv("20190306/data/S6_NOT_normalized_summarized_acetyl_abundance.txt")

#All peptides, normalized by protein expression.  Data is still normally distributed.
normalized_peptide_df <- tidy_df %>%
  group_by(Protein, Systematic_name, Peptide_id, Prot_PTM, Replicate, Time) %>%
  summarize(Intensity = mean(Intensity5)) %>%
  # group_by(Replicate, Time) %>%
  # mutate(Intensity = Intensity - median(Intensity)) %>%
  ungroup() %>%
  spread(Time, Intensity) %>%
  write_tsv("20190306/data/S6_normalized_summarized_peptide_abundance.txt")

#All peptides, normalized by protein expression.  Data is still normally distributed.
normalized_mod_df <- tidy_df %>%
  filter(grepl("[[:digit:]]Ac|Phospho| GG", Prot_PTM)) %>%
  group_by(Protein, Systematic_name, Prot_PTM, Replicate, Time) %>%
  summarize(Intensity = mean(Intensity5)) %>%
  # group_by(Replicate, Time) %>%
  # mutate(Intensity = Intensity - median(Intensity)) %>%
  ungroup() %>%
  spread(Time, Intensity) %>%
  write_tsv("20190306/data/S6_normalized_summarized_mod_abundance.txt")

#All peptides, NOT normalized by protein expression.  Data is still normally distributed.
peptide_df <- tidy_df %>%
  group_by(Protein, Systematic_name, Peptide_id, Prot_PTM, Replicate, Time) %>%
  summarize(Intensity = mean(Intensity4)) %>%
  # group_by(Replicate, Time) %>%
  # mutate(Intensity = Intensity - median(Intensity)) %>%
  # ungroup() %>%
  spread(Time, Intensity) %>%
  write_tsv("20190306/data/S6_NOT_normalized_summarized_peptide_abundance.txt")


#Used for mapping network functions onto proteins with acetylated peptides
Acetyl_map_ID <- tidy_df %>%
  filter(grepl("Acetyl", correct_PTM)) %>%
  group_by(Protein_id, Replicate) %>%
  summarize(Acetyl_marks = n_distinct(correct_PTM)) %>%
  filter(Acetyl_marks > 3) %>%
  left_join(Prot_table) %>%
  write_tsv("20190306/data/Acetyl_protein_table.txt")


#All peptides, normalized by protein expression.  Data is still normally distributed.
normalized_mod_df <- tidy_df %>%
  filter(grepl("[[:digit:]]Ac|Phospho| GG", Prot_PTM)) %>%
  group_by(Protein, Systematic_name, Prot_PTM, Replicate, Time) %>%
  summarize(Intensity = mean(Intensity3)) %>%
  # group_by(Replicate, Time) %>%
  # mutate(Intensity = Intensity - median(Intensity)) %>%
  ungroup() %>%
  spread(Time, Intensity) %>%
  write_tsv("20190306/data/S6_summarized_mod_abundance.txt")

Protein_summary_df <- tidy_df %>%
  filter(is.na(correct_PTM)) %>%
  group_by(Protein_id, Time, Replicate, Protein, Systematic_name, Gene_name) %>%
  summarize(Avg_Intensity4 = mean(Intensity3)) %>%
  spread(Time, Avg_Intensity4) %>%
  write_tsv("20190306/data/S6_summarized_unscaled_protein_abundance.txt")
```
