## Supplementary material for "Accurate and Sensitive Quantitation of the Dynamic Heat Shock Proteome using Tandem Mass Tags": File S1: S5_heatmaps_unscaled.R

```
library(tidyverse)
library(ggsci)
library(broom)
library(cowplot)
library(heatmaply)
library(ComplexHeatmap)

#tables
time_point_table <- read_tsv("bin/Time_point_table.txt")

df <- read_tsv("20190306/data/S6_summarized_mod_abundance.txt", guess_max = 100000)

df1 <- df %>%
  filter(grepl("[[:digit:]]Ac|Phospho| GG", Prot_PTM)) %>%
  mutate(Gene_name = gsub("\\_.*$", "", Prot_PTM)) %>%
  mutate(Prot_PTM = gsub("\\_\\, ", " ", Prot_PTM) %>%
           gsub("GG", "Ub", .) %>%
           gsub("Phospho", "Ph", .)) %>%
  unite(Gene_name_rep, Gene_name, Replicate, remove = FALSE) %>%
  select(Gene_name, Gene_name_rep, everything()) %>%
  filter(!Gene_name %in% c("SEC31", "SEY1", "HSP26")) %>%
  
  #Cleanup Prot_PTM names
  mutate(Prot_PTM = gsub("\\, \\,$", "", Prot_PTM) %>%
           gsub(" \\,", ",", .) %>%
           gsub("\\: Ub", "Ub", .) %>%
           gsub("\\: Ph", "Ph", .) %>%
           sub("\\, (?=s)", "_", ., perl = TRUE) %>%
           sub("\\, (?=t)", "_", ., perl = TRUE) %>%
           gsub("\\,$", "", .)
  )


prot_df <- read_tsv("20190306/data/S6_summarized_unscaled_protein_abundance.txt") %>%
  unite(Gene_name_rep, Gene_name, Replicate, remove = FALSE) %>%
  select(Gene_name, Gene_name_rep, everything()) %>%
  filter(Gene_name_rep %in% df1$Gene_name_rep)

df2 <- df1 %>%
  filter(Gene_name_rep %in% prot_df$Gene_name_rep,
         !grepl("Ox", Prot_PTM),
         !grepl("term", Prot_PTM)) %>%
  mutate(Prot_PTM = gsub("Ph\\_", "Ph, ", Prot_PTM) %>%
           gsub("Ub\\_", "Ub, ", .))


# Write to file -----------------------------------------------------------

df2 %>%
  filter(!grepl("\\, ", Prot_PTM)) %>%
  write_tsv("20190306/tables/20190329/Ac_Ph_Ub_single_mods.txt")

df2 %>%
  filter(grepl("\\, ", Prot_PTM)) %>%
  write_tsv("20190306/tables/20190329/Ac_Ph_Ub_multi_mods.txt")


prot_df %>%
  write_tsv("20190306/tables/20190329/Protein_abundances.txt")


# Complex heatmap ---------------------------------------------------------


# Phospho -----------------------------------------------------------------


pdf("20190306/plots/20190329/Ph_Heat_single.pdf", paper = "letter")
df2 %>%
  filter(grepl("Ph", Prot_PTM)) %>%
  filter(!grepl("\\, ", Prot_PTM)) %>%
  unite(a, Prot_PTM, Replicate) %>%
  select(a, `0`:`240`) %>%
  as.data.frame() %>%
  column_to_rownames("a") %>%
  as.matrix() %>%
  Heatmap(cluster_columns = FALSE,
          col = circlize::colorRamp2(c(-4,0,4), c("blue", "white", "red")),
          row_names_gp = gpar(fontsize = 6),
          heatmap_legend_param = list(title = "Normalized\nLog2 FC",
                                      title_gp = gpar(fontsize = 10)))
dev.off()


# Ubiquitin ---------------------------------------------------------------

pdf("20190306/plots/20190329/Ub_Heat_single.pdf", paper = "letter")
df2 %>%
  filter(grepl("Ub", Prot_PTM)) %>%
  filter(!grepl("\\, ", Prot_PTM)) %>%
  unite(a, Prot_PTM, Replicate) %>%
  select(a, `0`:`240`) %>%
  as.data.frame() %>%
  column_to_rownames("a") %>%
  as.matrix() %>%
  Heatmap(cluster_columns = FALSE,
          col = circlize::colorRamp2(c(-4,0,4), c("blue", "white", "red")),
          row_names_gp = gpar(fontsize = 8),
          heatmap_legend_param = list(title = "Normalized\nLog2 FC",
                                      title_gp = gpar(fontsize = 10)))
dev.off()


# Protein -----------------------------------------------------------------
pdf("20190306/plots/20190329/Protein_Heat.pdf", paper = "letter")
prot_df %>%
  select(Gene_name_rep, `0`:`240`) %>%
  as.data.frame() %>%
  column_to_rownames("Gene_name_rep") %>%
  as.matrix() %>%
  Heatmap(cluster_columns = FALSE,
          col = circlize::colorRamp2(c(-4,0,4), c("blue", "white", "red")),
          row_names_gp = gpar(fontsize = 6),
          heatmap_legend_param = list(title = "Normalized\nLog2 FC",
                                      title_gp = gpar(fontsize = 10)))
dev.off()


pdf("20190306/plots/20190329/Ubiquitin_protein_Heat.pdf", paper = "letter")
prot_df %>%
  filter(Gene_name %in% filter(df2, grepl("Ub", Prot_PTM))$Gene_name) %>%
  select(Gene_name_rep, `0`:`240`) %>%
  as.data.frame() %>%
  column_to_rownames("Gene_name_rep") %>%
  as.matrix() %>%
  Heatmap(cluster_columns = FALSE,
          col = circlize::colorRamp2(c(-4,0,4), c("blue", "white", "red")),
          row_names_gp = gpar(fontsize = 6),
          heatmap_legend_param = list(title = "Normalized\nLog2 FC",
                                      title_gp = gpar(fontsize = 10)))
dev.off()


pdf("20190306/plots/20190329/Phospho_protein_Heat.pdf", paper = "letter")
prot_df %>%
  filter(Gene_name %in% filter(df2, grepl("Ph", Prot_PTM))$Gene_name) %>%
  select(Gene_name_rep, `0`:`240`) %>%
  as.data.frame() %>%
  column_to_rownames("Gene_name_rep") %>%
  as.matrix() %>%
  Heatmap(cluster_columns = FALSE,
          col = circlize::colorRamp2(c(-4,0,4), c("blue", "white", "red")),
          row_names_gp = gpar(fontsize = 6),
          heatmap_legend_param = list(title = "Normalized\nLog2 FC",
                                      title_gp = gpar(fontsize = 10)))
dev.off()
```
