## Supplementary material for "Accurate and Sensitive Quantitation of the Dynamic Heat Shock Proteome using Tandem Mass Tags": File S1: Supplementary_Figure_1.R

```
library(tidyverse)
library(cowplot)
library(ggsci)

theme_set(theme_cowplot(font_size=12)) # reduce default font size

df<-read_tsv("data/CV_summary_timecourse.txt") %>%
  filter(SD > 0,
         Count < 10)


# Tidy ------------------------------------------------------------
Count_data <- df %>%
  group_by(Count, Replicate) %>%
  summarize(A = n())


(gg_violin <- df %>%
   filter(SD > 0) %>%
   ggplot() +
   geom_violin(aes(x = as.factor(Count), y=CV*100), fill="#56B4E9") +
   scale_y_continuous(trans="sqrt",
                      breaks = c(5, 25, 100, 200),
                      labels = c("5%", "25%", "100%", "200%")) +
    geom_text(data = Count_data, aes(label = A, x = as.factor(Count), y = 200)) +
    facet_wrap(~paste("Replicate",Replicate), nrow = 3) +
    labs(x="PSMs per peptide",y="CV%"))

ggsave("plots/Supplementary Figure 1.pdf", width = 8, height = 8, paper = "letter")
ggsave("plots/Supplementary Figure 1.tiff", width = 8, height = 8, dpi = 600, compression = "lzw")
```
