## Supplementary material for "Accurate and Sensitive Quantitation of the Dynamic Heat Shock Proteome using Tandem Mass Tags": File S5

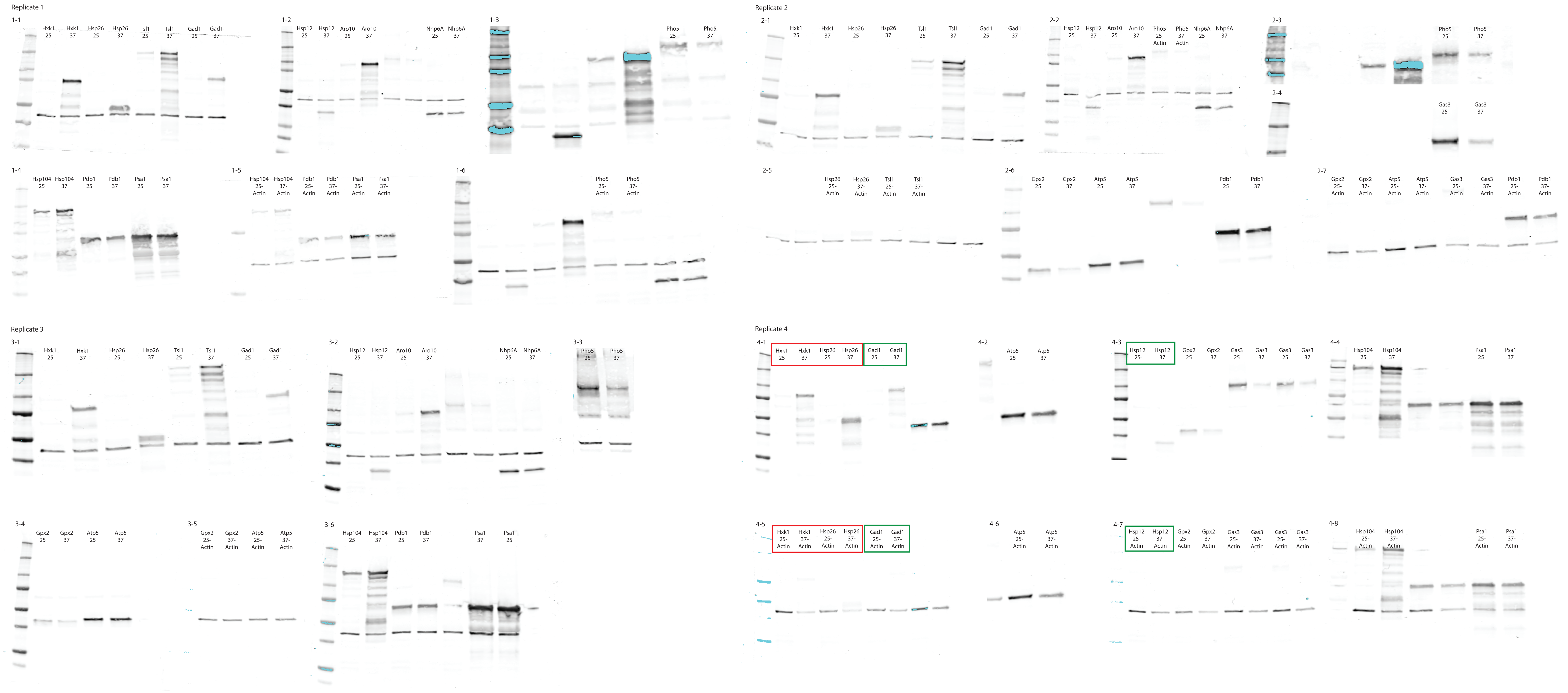


### Notes

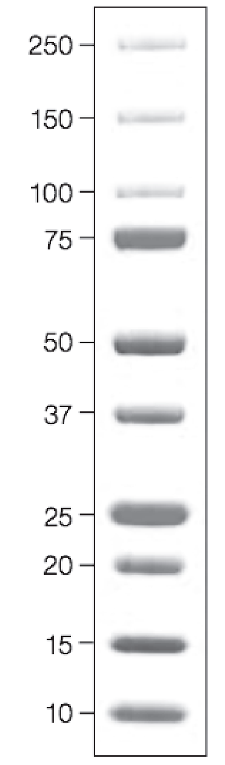


Ladder autofluoresces in the 700 channel only, and will only appear in images in which the 700 channel was on. (Same channel as TAP)

= Replicate 2 Redo

= Replicate 3 Redo

All unlabeled lanes were not used for calculations.

TAP and Actin quantification were perfoemd on the same gel unless otherwise noted.
